# α-Synuclein Triggers Intercellular Nanotubes Formation to Prevent Apoptosis in Astroglia by Promoting Stemness

**DOI:** 10.64898/2026.05.23.727344

**Authors:** Rachana Kashayap, Anirudh Sreenivas BK, Varshith MR, Rajashri Rameshwar Mundada, P Sreedevi, Shreshta Jain, Archanalakshmi Kambaru, Somasish Ghosh Dastidar, Sivaraman Padavattan, Vinay Kumar Rao, Ravi Manjithaya, Jiri Neuzil, Sangeeta Nath

## Abstract

Astrocytes play a significant role in neuroprotection by internalizing neurodegenerative aggregates and facilitating their degradation. Recent studies indicate that α-Synuclein (α-SYN) protofibrils promote the transfer of pathogenic aggregates and dysfunctional mitochondria between astroglia via tunneling nanotubes (TNTs), which enhances cell survival and resistance to apoptosis. However, the underlying mechanism of TNT-driven apoptosis resistance remains unclear. We find that α-SYN protofibrils induce aberrant mitochondria with decreased membrane potential (Ψm) and promote dynamic actin remodeling by relocating phosphorylated focal adhesion kinase (pFAK) to the nucleus, which triggers TNT formation in human astrocytoma cell lines and primary murine astrocytes. The important novel finding of this study is that pFAK in the nucleus co-localizes with Nanog, a crucial transcription factor for preserving stemness, and the interaction between pFAK and Nanog is critical for promoting p53 degradation via Mdm2-mediated ubiquitination and upregulating autophagy, thereby supporting the survival of astroglia exposed to toxic α-SYN protofibrils. ROCK inhibitor y-27632 also drives TNT-formation via pFAK translocation to the nucleus, colocalizes with Nanog, and enhances stemness-related gene expression. Inhibiting TNT with the actin depolymerizing agent cytochalasin-D prevents pFAK co-localization with Nanog in the nucleus and fails to protect cells from α-SYN-induced apoptosis. Nanog knockdown does not degrade p53 and hinders cell rescue from apoptosis. Furthermore, these transient TNTs transfer mitochondria to adjacent cells, potentially helping maintain metabolic stability. This study reveals that the TNT formation pathway promotes pFAK-Nanog interaction in the nucleus, leading to p53 degradation, which protects astroglia against α-SYN proteotoxicity and prevents apoptosis.

**Graphical Abstract:** 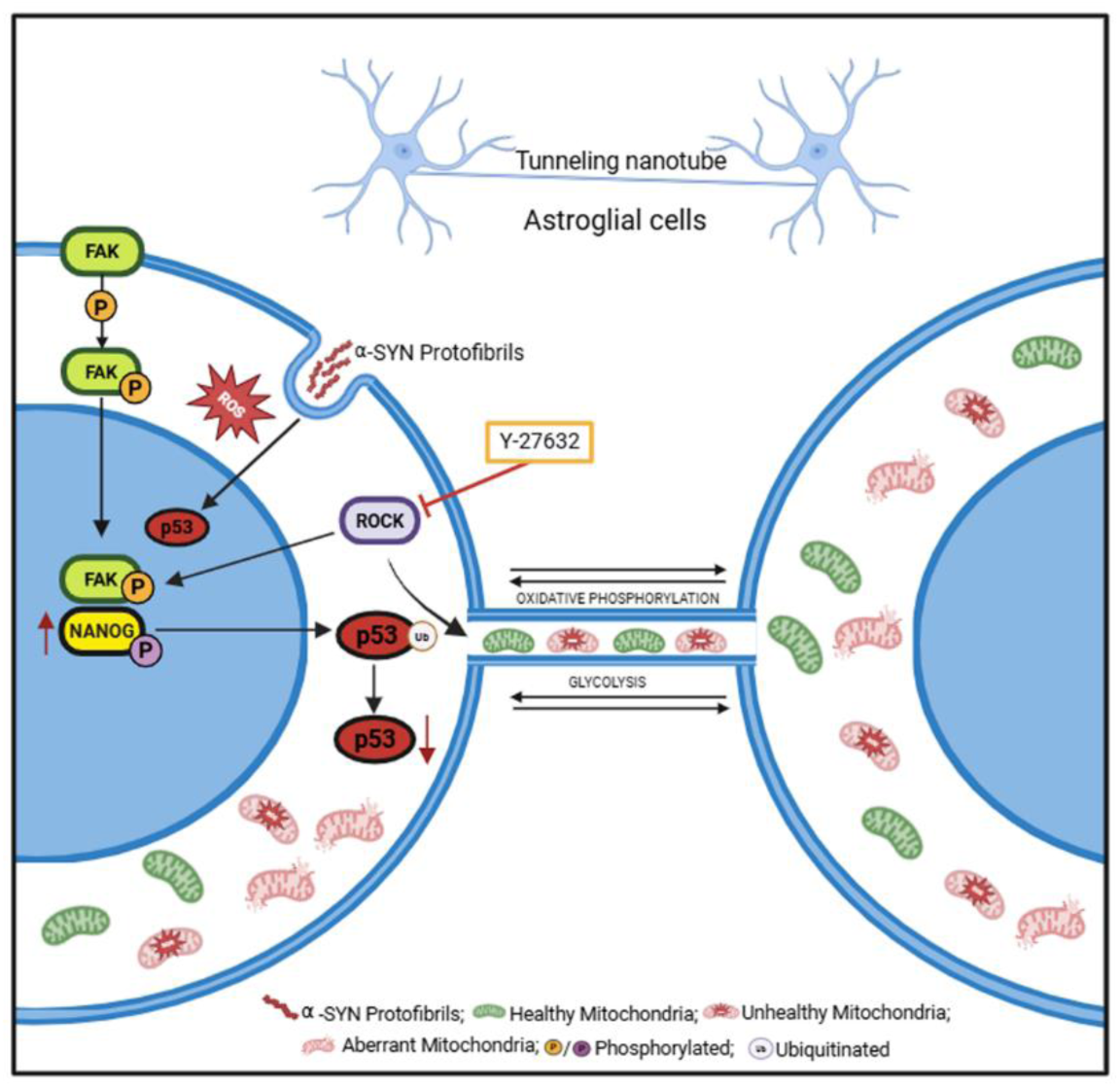

## Introduction

Neuroglial cells – microglia, astrocytes, and oligodendrocytes- play a crucial role in maintaining brain homeostasis and provide protection against proteinopathies seen in neurodegenerative diseases like Alzheimer’s Disease (AD) and Parkinson’s Disease (PD) [1], [2]. Intercellular communications between neurons and glial cells contribute to the spread of PD pathology and neuroinflammation [3], [4], [5]. Various modes of intercellular communication, including gap-junctions, exosomes, paracrine secretions, and the recently identified tunneling nanotubes (TNTs), are essential for neuron-glia interactions [6], [7], [8]. Studies indicate that microglia and astrocytes enhance TNT-mediated communication when exposed to extracellular aggregates of amyloid-β and α-SYN. This interaction aids in the intercellular spread of the toxic load transfer between glial cells, which helps reduce pathogenic stress and promotes rapid clearance of neurodegenerative aggregates [9], [10], [11], [12].

Recent studies have shown TNTs to be narrow-bore (diameter around 50-700 nm), open-ended structures with intercellular cytoskeletal-actin continuity (up to 300 µM) for direct transfer of cytoplasmic content, including organelles [13], aggregates of neurodegenerative proteins [14] and pathogens [15]. TNTs primarily form in response to various types of cellular or oxidative stress, but their duration is transient. Several recent studies have highlighted their protective role, particularly in stressed cells, by mitigating apoptosis and promoting cell survival [10], [14], [16], [17], [18] The TNTs enable the sharing of proteotoxic burdens, enhancing communication between neurons, microglia, and astrocytes, which is crucial for maintaining neuronal health and mitochondrial balance. On similar lines, our previous study also revealed that TNT formation precedes clearance of α-SYN-induced organelle (lysosomes and mitochondria) toxicities, alleviating ROS levels and reversing premature cellular senescence in the primary astrocytes and astroglia cells [12]. Microglia also employ a similar mechanism of TNT formation and intercellular transfer of toxic burden to rescue themselves from α-SYN-induced toxicity [7], [19]. Cells containing mitochondria devoid of mtDNA can regain their function by acquiring functional mitochondria through TNTs from nearby stromal cells, aiding in cells’ energy restoration [20], [21]. Additionally, studies have shown that the intercellularly transferred aberrant mitochondria with accumulated ROS promote cell proliferation [12], [22].

Similar mechanisms of TNT-mediated mitochondrial transfer have also been observed in astrocytoma/glioblastoma [23], [24], [25] - the most common primary tumour of the central nervous system (CNS), mainly originating from astrocytes [26]. Various conditions, including oxidative stress, radiotherapy, and chemotherapy, promote the formation of TNTs, which could drive the transformation of astrocytes into cancer stem-like cells [23], [27]. It was also reported that ROCK inhibition promotes expression of stem-like phenotypic genes during the *in vitro* expansion of astrocytoma/glioblastoma cells [28]. These cells are highly dynamic and make the tumour resistant to radiation and chemotherapy [29], [30], [31], [32] TNTs can help cells resist apoptosis by allowing stressed cells to receive functional mitochondria from healthy neighbors, restoring their function and preventing cell death. TNT-mediated mitochondrial transfer has also been observed within 3D-organoid models from patient-derived glioblastoma stem cells [33]. Research suggests that P53, a protein that induces apoptosis, may be involved in TNT biogenesis, but its role is complex and context-dependent [17], [34], [35]

Proteotoxic stress within cells leads to mitochondrial damage and elevated ROS, prompting a metabolic switch from oxidative phosphorylation to glycolysis, akin to the Warburg effect observed in cancers [36]. The proteotoxic stress to the cells that ‘inherit’ anti-apoptotic properties makes them more likely to survive, enhancing the TNT formation [17]. However, the mechanisms by which α-SYN protofibril-treated astrocytes adapt to stem-like cells—promoting TNTs to mitigate mitochondrial toxicity and improve cell survival preventing apoptosis —are not fully understood.

This study shows that α-SYN-treated astrocytes and astroglia induce TNTs, facilitating mitochondrial transfer and recovery from oxidative stress. RNA sequencing revealed upregulation of stemness markers and a metabolic shift to glycolysis as a part of the cell’s survival strategy upon α-SYN treatment. This can be attributed to the reduction in aberrant mitochondria following TNT-mediated clearance via cell-to-cell transfer. α-SYN and ROCK inhibitor-treated glial cells express increased levels of the stemness marker – Nanog, in response to nuclear translocation of pFAK during α-SYN protofibrils-induced TNT-formation. The interaction of pFAK and Nanog, led to p53 degradation by promoting its ubiquitination, which is vital for the survival of α-SYN protofibrils-treated astroglia. These findings highlight the role of transient nuclear translocation of pFAK during TNT formation in promoting survival of α-SYN protofibrils-treated astroglia, preventing p53-mediated apoptosis, and enhancing the expression of pluripotency-associated transcription factors.

## Material and methods

The work has been reported in line with the ARRIVE guidelines 2.0.

### Mice

Six to eight-weeks-old C57BL/6J mice, regardless of sex, were obtained from Jackson Laboratories. They were kept under a 12-hour dark-light cycle at a relative humidity of 50–60% and a temperature of 25°C. Both male and female mice were included in the study without bias.

### Primary astrocyte culture

Six to eight-weeks-old C57BL/6J mice of either sex were euthanized using CO₂ asphyxiation, and the cerebral cortices were carefully dissected from their brains. The tissue samples were thoroughly washed with cold HBSS, and meningeal layers were removed by rolling the tissues over sterile Whatman filter paper. The cleaned tissues were triturated using a 1 mL pipette in the presence of a digestion buffer containing 0.25 mg/mL collagenase, 0.25% trypsin-EDTA solution (Gibco, #25200-072), 0.25% FBS (Gibco, #1600004), and 1x PBS. The tissue suspension was incubated at 37°C for 30 minutes to facilitate dissociation. Following incubation, the mixture was centrifuged at 1500 rpm for 5 minutes, and the resulting cell pellet was resuspended and plated in MEM supplemented with 20% FBS on Corning T25 TC-treated culture dishes. The medium was replaced with fresh MEM containing 20% FBS the following day, and the cells were maintained under these conditions for 14 days. Subsequently, the cells were trypsinised and re-plated as required for experiments.

### Cell lines and culture medium

U-87 MG and U251 cell lines, which are derived from astrocytoma-glioblastoma cancer, were obtained courtesy of Prof. Kumaravel Somasundaram at the Indian Institute of Science, Bangalore, India [37]. The original source of U251 cell line was from the European Collection of Authenticated Cell Cultures (ECACC, # 09063001). The Certificate of general cell collection procedure and approvals are available at https://www.culturecollections.org.uk/products/celllines/generalcell/detail.jsp?refId=09063001&collection=ecacc_gc. The U87 cell line original source is ECACC (#89081402). The Certificate for all approvals is available at https://www.culturecollections.org.uk/products/celllines/generalcell/detail.jsp?refId=89081402&collection=ecacc_gc. These cells were tested for mycoplasma contamination (using the Sigma-Aldrich #M7006 kit). Cells were cultured in Dulbecco’s Modified Eagle Medium (Gibco #2120395), supplemented with 10% Fetal Bovine Serum (Gibco #1600004) and 1% PSN (Penicillin-Streptomycin-Neomycin Mixture; Thermo Fisher Scientific #15640055) at 37°C in a 5% CO_2_ humidified incubator. The glioblastoma cell lines U251 and U87 both exhibit astroglial characteristics and express the Glial Fibrillary Acidic Protein (GFAP) marker.

### Preparation of α-SYN protofibrils and treatment conditions

α-SYN wild-type (Addgene ID #36046) construct was overexpressed in E.coli and purified [38]. α-SYN protofibrils were prepared by incubating 1 μM of protein with 0.65% 4-hydroxynonenal (HNE) (10mg/ml stock) (Sigma-Aldrich #393204-1MG) at 37 °C for 7 days with moderate shaking, as described in Raghavan et al. 2024. The protofibrils were subjected to lyophilization and subsequently stored at −80 °C following characterization by transmission electron microscopy (TEM) until the commencement of the experiment. TEM samples were prepared similarly as described previously[12].

The experimental setup involved treating primary astrocytes, U-87 MG, and U251 cells with 1 µM α-SYN protofibrils at 37°C, 5% CO₂ for durations of 3, 6, 12, and 24 hours. All cells, including untreated controls, were seeded simultaneously to ensure uniformity. After 24 hours of seeding, treatments were started once the cells had adhered and looked healthy. The treatment process followed a reverse sequence of time points to allow simultaneous analysis across all groups. Specifically, cells designated for the 24-hour treatment were exposed to α-SYN protofibrils first, followed by the 12-hour, 6-hour, and 3-hour groups. Untreated control cells were maintained under identical conditions throughout, serving as a baseline for comparison across all treatment durations. All subsequent experiments were conducted simultaneously for treated and control groups to maintain consistency in experimental conditions.

### RNA sequence analysis

Raw sequence reads were subjected to FastQC (v0.12.1) to assess read quality and reads were further trimmed using Trimmomatic (v0.39). The trimmed reads were aligned to the reference genome using HISAT2 (v2.2.1) in a stand-alone custom analysis pipeline. The aligned reads were quantified to generate raw counts using FeatureCounts (v2.0.8). Raw counts were then normalized through the median of ratios method, accounting for the sequencing depth and RNA composition bias; further differential expression analysis was performed using DESeq 2 (v1.40.2) by modelling the counts using a negative binomial distribution and applying a Wald test to determine the statistical significance. Furthermore, p-values were adjusted for downstream analysis using the Benjamini-Hochberg method to control the false discovery rate (FDR). Differentially expressed genes were further categorized by fold-change ≥ 1.5 and FDR-adjusted p-value of < 0.05. Gene Ontology (GO) was performed using gProfiler with default parameters, while the Gene Set Enrichment Analysis (GSEA) database was used for pathway enrichment analysis. The volcano plot, cluster heatmap, and pathway categorization were all plotted using R (v4.4.2) using ggplot2 (v3.5.1), enhanced volcano (v1.24.0), and dplyr (v1.1.4) packages.

### Immunocytochemistry

For immunofluorescence staining, cells (astrocytes, U87 and U251 cells) were seeded onto coverslips pre-coated with Poly-D-Lysine or in glass-bottom 35 mm dishes (Cellvis, D35-14-1.5-N) at 10,000 cells. Cells were treated with α-SYN protofibrils (1μM) and Y-27632 (5μM) for durations of 3, 6, 12, and 24 hours. The experimental setup with inhibitors involved 5 μM Y- 27632 [ROCK inhibitor (Sigma-Aldrich #Y0503)], 0.5 μM Cytochalasin-D (Thermo Fisher Scientific #PHZ1063), 5mM *N*-Acetyl-L-cysteine [NAC (Sigma-Aldrich #A9165)] with and without α-SYN protofibrils (1μM) and Y-27632 (5μM) for durations of 3, 6, 12, and 24 hours before proceeding with subsequent experiments.

After treatment, cells were washed with 1x PBS and fixed with 4% PFA for 20 minutes. The cells were then permeabilized and blocked in an incubation buffer containing 1 mg/ml saponin and 5% FBS for 20 minutes at room temperature. Cells were then incubated overnight at 4°C with the specified primary antibodies and secondary fluorescently labeled antibodies. Then, ProLong Gold antifade reagent with DAPI (Invitrogen P36941) was used as a counterstain.

Primary antibodies Ki67 anti-rabbit (Millipore #B9260) as cell proliferation marker; FAK Rabbit (CST #3285T) and phospho-FAK (Tyr397) (Invitrogen #44-624G) as total and active FAK marker; CD133 (Invitrogen #14133182) as cancer stem cell marker; Nanog (Invitrogen, #MA1-017) and Sox2 (Cloud-Clone#PAA406Hu01) as stem-like progenitor markers; GFAP (Cloud-Clone, #PAA068Hu01) as astrocytic marker; Phalloidin conjugated with iFlour555 (Abcam #176756) were used to stain actin in TNTs. Secondary antibodies Alexa- 488 goat anti-rabbit (#A-11070; Invitrogen), Alexa- 555 anti-mouse (#A-1413312; Invitrogen), Alexa-488anti-mouseIgG(H+L) (#A-11059; Invitrogen), Alexa-555anti-rabbit (#A-21428; Invitrogen) were used respectively following the above-mentioned protocol. All primary antibodies were used at a dilution of 1:300, secondary antibodies at 1:500, and phalloidin at 1:700.

Image acquisition was performed using either a laser confocal microscope (Zeiss LSM880, Carl Zeiss, Germany) or a fluorescence microscope (Olympus IX73). The captured images were subsequently processed and analysed using ImageJ software.

### Nuclear fractionation

U-87 MG cells were seeded at 200,000 cells per well in 60 mm culture dishes and treated with 1 µM α-SYN protofibrils (1μM) for durations of 3, 6, 12, and 24 hours. Following treatment, cells were harvested using 0.25% trypsin-EDTA solution (Gibco, #25200-072, Canada origin). The cell pellet was resuspended in cold cytoplasmic extract (CE) buffer (added at five times the pellet volume) and incubated on ice for 3 minutes. The composition of CE buffer is NP40 detergent (0.075 %) with EDTA, DTT, and PMSF each 1 mM in 10 mM HEPES buffer (pH 7.6) supplemented with KCl (60 mM). The suspension was centrifuged at 1500 rpm for 5 minutes at 4°C, and the supernatant, representing the cytoplasmic extract, was collected into a prechilled tube. The remaining nuclear pellet was washed with CE buffer (without detergent), centrifuged at 1500 rpm for 5 minutes at 4°C, and resuspended in nuclear extraction (NE) buffer (added at two times the pellet volume). The composition of NE buffer is glycerol (25 %) with EDTA (0.2 mM), MgCl2 (1.5 mM), and PMSF (1 mM) in 20 mM Tris-Cl buffer (pH 8) supplemented with NaCl (420 mM). The mixture was incubated on ice for 10 minutes, followed by centrifugation at 1500 rpm for 5 minutes at 4°C. The resulting supernatant, containing the nuclear extract, was collected into a prechilled tube. Both cytoplasmic and nuclear extracts were stored at −80°C for further analysis.

### Immunoprecipitation

U-87 MG cells were seeded at 2,200,000 cells per 100 mm culture dish and treated with 1 µM α-SYN protofibrils (1μM) for 24 hours. To inhibit proteasomal degradation, cells were pretreated with 10 μM MG-132 (Merk, #133407-82-6) for 5 hours prior to lysis. Following treatment, cells were lysed using RIPA buffer (50 mM Tris-HCl, pH 8.0, 150 mM NaCl, 1% NP-40, 0.5% sodium deoxycholate, 0.1% SDS, 1× protease inhibitor, and 25 mM NEM). The cell lysates were incubated overnight at 4°C with anti-P53 (Invitrogen, #MA512557) antibody. Dyna Green magnetic beads (Invitrogen, #80104G) were pre-washed with 400 µl RIPA buffer and then added to the antibody–lysate mixture (20 µl per sample). Samples were rotated for 2 hours at 4°C, after which the beads were collected and washed with RIPA buffer. Following the final wash, the beads were resuspended in 50 µl of 1× SDS loading dye and boiled at 98°C to elute bound proteins.

### Western blot

Western blot analysis was conducted after normalizing protein concentrations across samples. The membranes were probed with primary antibodies, including H3 (Invitrogen, #865R2), Nanog (Invitrogen, #MA1-017), phospho-Nanog (Ser71) (Invitrogen, #PA5-13078), P53 (Invitrogen, #MA512557), p21 Waf1/Cip1 (CST #2947), and Ubiquitin (Invitrogen, #14-6078-82), Caspase-3 (Cloud clone, #MAA626Hu01), GAPDH (Cloud clone, #CAB932Hu22), beta-3 Tubulin (Invitrogen, #22833). Primary antibody dilutions are 1:1000. Secondary antibodies used were Goat anti-Mouse (H+L) (Invitrogen, #32430, 1:1500 dilution) and Goat anti-Rabbit (H+L) (Invitrogen, #32460, 1:1500 dilution). The blot signals were developed using ECL solution (SuperSignal West Femto Trial Kit, Invitrogen, #34094). Band intensities were quantified via densitometry using the gel analyzer plugin in Fiji software.

### Caspase-3 activation analysis

Caspase-3 activation was assessed by Western blot analysis of pro-Caspase-3 (∼35 kDa) and cleaved Caspase-3 (∼15 kDa). Band intensities were quantified using ImageJ. Both pro and cleaved Caspase-3 levels were normalized to GAPDH followed by assessing the ratio of cleaved to pro-Caspase-3 to determine relative Caspase-3 activation.

### Live cell imaging by DiD Vybrant membrane dye

U-87 MG cells were seeded at 80,000 cells per well in a 35 mm dish. Treatments were conducted as previously described using α-SYN protofibrils (1μM), NAC (5mM), and cytochalasin-D (Cyto-D) (0.5μM) (Thermo Fisher Scientific #PHZ1063) for durations of 3, 6, 12, and 24 hours. Following treatment, cell membranes were labeled with the DiD Vybrant dye (Invitrogen #V22887).

The dye was prepared by diluting it 1:200 in DMEM (phenol red-free) supplemented with 10% FBS and incubating the cells at 37°C for 20 minutes. After incubation, the cells were washed with DMEM for 10 minutes to remove excess dye. Live-cell imaging was then performed using a fluorescence microscope (Olympus IX73) to visualize and assess the presence of thin membrane structures resembling TNTs.

### Mitochondria transfer in co-culture

U-87 MG cells were transiently transfected with pLV-mitoDsRed (Addgene#44386; was a gift from Dr.Pantelis Tsoulfas’s lab) and co-cultured with an equal population of mEGFP-lifeact-7 (Addgene#58470; was a gift from Michael Davidson) transfected cells. The transfections were performed using lipofectamine 3000 (Invitrogen #44386). Both the transfected cells were mixed at a density of 60,000 cells in glass-bottom 35mm dishes (Cellvis, #D35-14-1.5-N) and treated with Y-27632 (5μM) for durations of 3, 6, 12, and 24 hours. The cells were then fixed with 4 % PFA before imaging using a confocal microscopy.

### Transfection and transduction of shRNA-Nanog

HEK 293T cells were seeded at 2,200,000 cells per 100 mm culture dish. The transfection reagent, FuGENE® HD, was prepared by diluting it in OptiMEM at a 1:4 ratio relative to the plasmid concentration. The mixture was briefly vortexed and incubated for 5 minutes at room temperature. To the prepared transfection mix, 4-10 μg of viral packaging plasmids, including the transfer plasmid [pLKO.1] integrated with shNanog [Sigma Aldrich, SHCLNG TRCN0000004884], envelope plasmid [pMD2.G (addgene id: #12259)], and packaging plasmid [psPAX2 (addgene id: #12260)], were added. The mixture was incubated for 30 minutes at room temperature before being applied to the HEK 293T cells. The cells were incubated at 37°C in a CO₂ incubator. The culture supernatant, containing viral particles, was harvested at 48 and 72 hours post-transfection. The collected supernatant was centrifuged at 4500 rpm for 30 minutes at 4°C to concentrate the viral particles. These concentrated particles were then used to transduce recipient U-87 MG cells. After 24 hours, puromycin (1 mg/ml; HiMedia) was added to the cultures for selection of successfully transduced cells. The selected cells were subsequently used for further experiments.

### Preparation of spheroid cultures using the hanging drop method

Primary astrocytes, wildtype and shRNA-Nanog knockdown U-87 MG cells were plated at 80,000 cells per well in a 24-well plate and treated with 1 µM α-SYN protofibrils, 5 µM Y-27632, 0.5 µM Cyto-D, 5 µM Y-27632 + 1 µM α-SYN protofibrils, and 0.5 µM Cyto-D + 1 µM α-SYN protofibrils for durations of 3, 6, 12, and 24 hours. Post-treatment, cells were detached using 0.25% trypsin-EDTA solution (Gibco, #25200-072, Canada origin) and cultured using the hanging drop method. A 20 µL drop of cell suspension was placed on the lid of a 100 mm sterile non-adherent plate, with 1x PBS added to the bottom of the plate to maintain humidity and support spheroid formation. After 24 hours of culture, spheroids were imaged, and the size of the spheroids were analysed.

### Mitochondrial membrane potential

Mitochondrial membrane potential was assessed using tetramethylrhodamine, ethyl ester, perchlorate (TMRE) (Invitrogen #T669) staining. When the membrane potential is high, TMRE accumulates in mitochondria, emitting a red-orange fluorescence (λ_ex/em_: 549 / 574 nm). U-87 MG cells were seeded at 80,000 cells per well in a 35 mm dish. Treatments were performed as previously described using α-SYN protofibrils for durations of 3, 6, 12, and 24 hours. After treatment, cells were incubated with 100 nM TMRE at 37°C for 30 minutes. Fluorescence intensity was measured using an Olympus CKX53 microscope with an excitation wavelength of 540 nm and an emission wavelength of 560 ± 20 nm.

### Estimation of ROS by flowcytometry

U-87 MG cells were seeded at 100.000 cells per well in a 24-well plate and treated with 5 µM Y-27632 for durations of 3, 6, 12, and 24 hours. Following treatment, the cells were detached using 0.25% trypsin-EDTA solution (Gibco, #25200-072, Canada origin). The harvested cell pellets were resuspended in DMEM supplemented with 10% FBS and 20 µM 2’-7’-Dichlorodifluorescein diacetate (DCFDA) for reactive oxygen species (ROS) detection. The cells were incubated with DCFDA for 30 minutes at 37°C, after which they were washed with 1x PBS to remove excess dye. ROS levels were assessed by measuring the fluorescence of DCFDA-stained cells using a flow cytometer (BD LSR II) at an excitation/emission wavelength of 488/520 nm. Data analysis was conducted using FCS Express software to quantify fluorescence intensity.

### β-galactosidase activity assay

U-87 MG cells were seeded at 80,000 cells per well in a 35 mm dish. Subsequently, cells were treated with α- SYN protofibrils (1μM) for durations of 3, 6, 12, and 24 hours. Post treatment, β-galactosidase activity was detected using β-Galactosidase Staining Kit (Cell Biolabs, Inc #AKR-100). Following the staining, brightfield images were captured using a coloured camera and analysed for β-galactosidase-positive cells, a hallmark of cellular senescence.

### TNT characterisation

Cells were stained using phalloidin to identify actin conduits using confocal images. The images were analysed using Fiji a Java-based image processing software developed at the National Institutes of Health (NIH) and the Laboratory for Optical and Computational Instrumentation (LOCI). TNTs were distinguished by their unique ability to hover without attaching to the substratum, and identified from reconstructed 3D volume view using plugin of Fiji as described earlier in [39]. TNTs were manually counted and plotted as the ratio of the number of TNTs to the number of cells per field.

### Fluorescence intensity analysis

The expression levels of Ki67, FAK, pFAK, Sox2, Nanog, and CD133, p53 and propidium iodide were quantified by measuring the intensity of labeled proteins per cell in the images for each condition. Intensity analysis was performed using Fiji, with regions of interest (ROIs) defined and selected using the ROI-plugin.

### Area analysis

Spheroid area was quantified using Fiji, where regions of interest (ROIs) were defined and selected with the ROI-plugin, followed by measurement using the Measure option.

### Statistics and reproducibility

In the bar graphs, data are presented as mean ± standard deviation (SD). Individual data points are shown as dots. Each figure legend includes key details such as mean values, standard deviations, the number of images used for the analysis, biological or technical replicates, statistical tests used, and any corrections applied. For Western blot analysis, relative protein abundance was determined by measuring signal intensity relative to a loading control. To ensure comparability across different blots, data were normalized by dividing each value by the sum of all data points on the same membrane. To validate the significance of the analysed data, two-tailed Student’s *t* test was used to compare two different groups; a one-way analysis of variance (ANOVA) was used to compare more than two groups. For data with two independent variables, a two-way ANOVA was used. The confidence interval was 95%. \**P* < 0.05, \*\**P* < 0.01, \*\*\**P* < 0.001, and \*\*\*\**P* < 0.0001.

## Results

### TNTs regulates mitochondrial fate in the α-SYN protofibrils treated toxic astroglia cells

Our recent work demonstrated that α-SYN protofibrils induce proteotoxic stress, promoting transient biogenesis of TNTs in astrocytic lineage cells, such as U87-MG, U251 astrocytoma cell lines, and primary murine astrocytes in the initial 3 and 6 hours of the treatment [12]. The transient formation of TNTs is closely linked to a decrease in mitochondrial membrane potential, as assessed by the JC1 dye [40]. In this study, we demonstrate that formation of transient TNTs (at early times, 3 and 6 hours) in primary astrocytes (Figure 1A-B) is associated with aberrant mitochondria exhibiting decreased mass and lower mitochondrial potential, as evaluated by TMRE dye (Figure 1C-D). α-SYN protofibrils-treated cells featuring multiple TNTs at early time points undergo adaptation to recover the normal state characterized by polarized mitochondria by 24 hours (Figure 1C, D).

**Figure 1:**
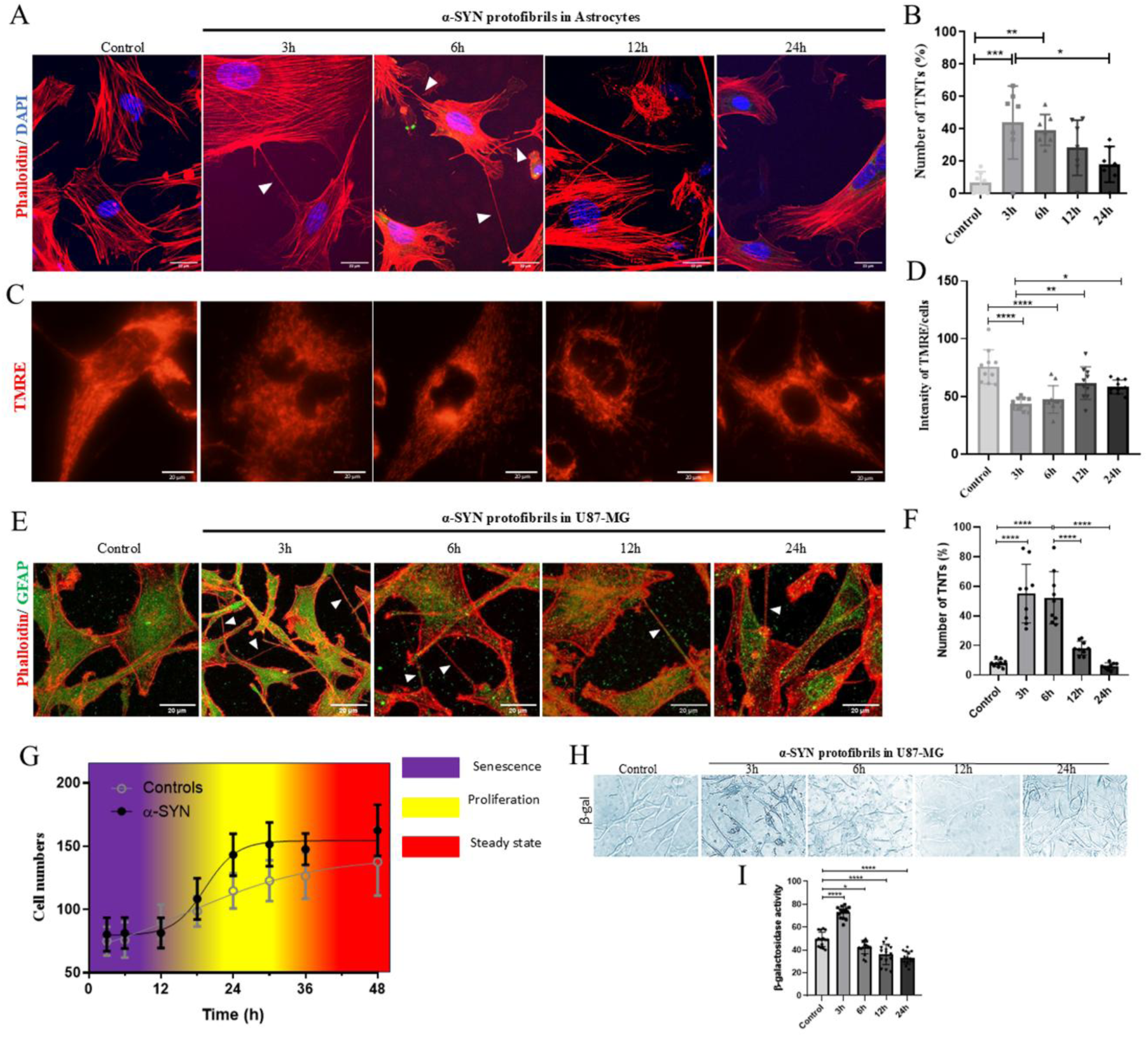
Effect of α-SYN protofibrils (1 μM) on TNT-biogenesis in correlation to mitochondrial toxicity and proliferation in astroglia. A) Transient increase of TNT biogenesis (white arrows) in the primary astrocytes treated with α-SYN-protofibrils at 3 and 6 h, compared to the control. B) Quantification of the percentage of TNTs in astrocytes from 6 image frames containing >15 cells for each set. C)Mitochondrial membrane potential measurements using TMRE dye in α-SYN-treated astrocytes. Intensities from the cells were counted from randomly taken 4 images containing >10 cells for each set. D) Quantification of TMRE intensities as an indicator of Ѱ of mitochondria. E) Transient increase of TNT biogenesis (white arrows) in the U87 MG cells (positive for GFAP) treated with α-SYN-protofibrils at 3 and 6 h, compared to the control. F) Quantification of the percentage of TNTs in U87 MG cells from 6 image frames containing >15 cells for each set. G)Growth curve of control and 1µM α-SYN protofibrils treated U-87 MG cells, readings were taken across 3 h - 48 h and the cells counted manually (counted from randomly taken > 10 images for each set). H) β-Galactosidase activity in U87 MG cells treated with 1µM α-SYN protofibrils. I) Quantification of β-Galactosidase activity. Data are expressed as mean ± SD, *** p ≤ 0.001. Statistics were analysed using two-way ANOVA. N=3.

The U-87 MG cells exhibit a similar trend of transient TNTs formation at early times 3 and 6 hours (Figure 1E, F). U-87 MG astrocytoma cells, though transformed tumour cells, are known to display key astrocyte traits, such as GFAP expression and radial branching. We validated GFAP expression in U-87 MG cells, and the results show that TNTs in these cells are GFAP-positive (Figure 1E). A similar phenomenon of mitochondrial membrane potential loss was observed for U-87 MG astrocytoma cells treated with α-SYN protofibrils using a mitotracker dye (Figure S1A). Staining with mitotracker allowed for discerning the fragmentation pattern of mitochondria with its reduced intensity at 3 and 6 hours after treatment of U87-MG cells compared to control cells and to the treated cells at 24 hours post-treatment (Figure S1A). Cells treated with α-SYN protofibrils exhibit numerous TNTs 3 hours after treatment, allowing for mitochondrial movement (Figure S1B). Protofibrils of α-SYN were validated for each batch using TEM, one of which is represented in Figure S1C. The experiment with HNE was performed as a vehicle control to confirm that TNT formation is solely caused by α-SYN protofibrils, with no effect from the vehicle control (Figure S1D,E).

Nevertheless, to investigate how cells are affected by treatment with α-SYN protofibrils, we conducted cell growth curves and compared them to the growth of control cells. Cells exhibit a transient initial growth arrest (3-6 hours) after α-SYN treatment. Later, the cells recovering from α-SYN toxicity after 24 hours exhibited higher rate of proliferation compared to control U87 MG astrocytoma cells (Figure 1G). The early transient stress-induced growth arrest resembles premature senescence-like characteristics, as evidenced by positive β-galactosidase activity at 3 and 6 hours (Figure 1H,I). This is consistent with our earlier study, which revealed transient early senescence-like phenotypic characteristics, as validated by p21 expression and β-galactosidase activity upon α-SYN exposure [12].

### α-SYN-induced proteotoxic stress initiates transcription level change in gene expression to safeguard astroglia cells against mitochondrial toxicity

Previous studies, including our own, demonstrated that astrocytic lineage cells (astrocytoma/glioblastoma cell line and primary astrocytes, termed as astroglia) efficiently degrade toxic α-SYN aggregates, promote TNT formation, and aid in recovery by encouraging proliferation [9], [10], [11], [12]. This study examines how astrocytic lineage cancer cells and primary astrocytes withstand toxic neurodegenerative aggregates and both share survival traits similar to those of astrocytoma/glioblastoma. Thus, RNA sequencing data analysis was conducted to comprehend alterations in gene expression resulting from α-SYN-induced proteotoxic stress in U87-MG cells. We found that it triggered metabolic reprogramming in the cells by shifting from oxidative phosphorylation to glycolysis during the early period (3 hours) (Figure 2A-D). This activation led to increased expression of stem-like progenitor markers, presumably as a cellular survival strategy. The enhanced oxidative stress response resulting from exposure to α-SYN protofibrils led to a significantly high-level increase in mitochondrial toxicity, depicted by amplification of processes such as mitochondrial fission with concomitant downregulation of the mitophagy pathway (Figure 2C). These adaptive changes help in rescuing the cells from oxidative stress at later times (24 hours) by upregulating mitophagy and downregulating mitochondrial fission, and reversing the glycolysis pathway (Figure 2D). Upregulation of stemness markers occurs both at 3 and 24 hours after the α-SYN protofibril treatment (Figure 2A-D).

**Figure 2:**
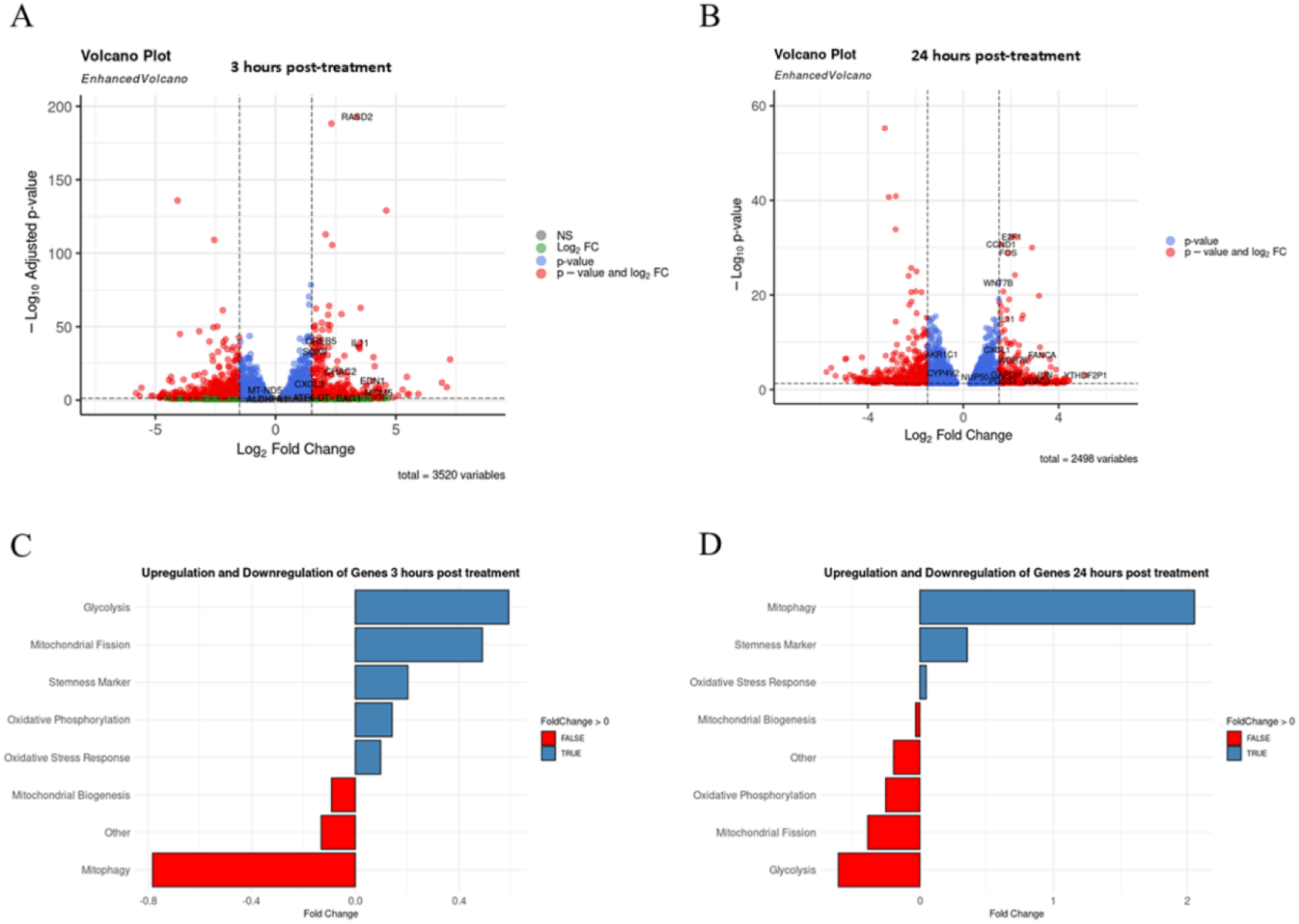
RNA sequence data to show pathway upregulation and downregulation upon α-SYN protofibrils treatment. The volcano plot, and pathway categorization was plotted from RNA sequence data. volcano plot of A)3 h – and B) 24 h-post-treated U87-MG astroglia cells with α-SYN protofibrils. C-D) Pathway categorization from upregulation and downregulation of genes A)3 h – and B) 24 h-post-treated U87-MG astroglia cells with α-SYN protofibrils.

Furthermore, the volcano plot highlights upregulated genes linked to oxidative stress-induced senescence: RASD2, IL11, CHAC2 and MCM5 genes evaluated at the 3-hour time point after α-SYN protofibril treatment (Figure 2A). α-SYN toxicity also inhibits the expression of mitophagy gene MAP1LC3Bd during early time points. The volcano plot also indicated upregulation of the genes SOX9 and EDN1, which are associated with stemness, as well as the genes CREB5 and CXCL3, which relate to cancer cell response to drug resistance. Additionally, the analysis revealed upregulation of genes related to glycolysis, and downregulation of genes in the oxidative phosphorylation (Figure 2A).

Adaptation of astroglia cells to proteotoxic stress involves transition to stemness, as indicated by involvement of genes such as E2F1, FOS, CXCL1, JUN, and VDAC3, which are linked to the maintenance of cancer stem-like cells, along with WNT7B, IL11 and YTHDF2PI at later time of 24 hours (Figure 2B). At 24-hour time point after exposure to α-SYN, cells exhibit reduced expression of glycolysis (GAPDH) and oxidative phosphorylation (AKR1C1, CYP4V2) genes compared to control cells (Figure 2B). These findings indicate that proteotoxic stress induced by α-SYN initiates cellular adaptation within the initial period, resulting in a shift in metabolic dependence from oxidative phosphorylation to glycolysis. This activation culminates in the altered expression of stem-like progenitor markers and drug-resistance genes, reflecting a strategy employed by the cells to shift toward enhanced survival.

### Nuclear translocation of pFAK upregulates expression of stemness markers

We next studied cell proliferation and enhanced stemness in astrocyte-origin cancer cell lines, exposed to α-SYN protofibrils. We conducted experiments using the U-87 MG and U251 astrocytoma cell lines to examine overall effects on astrocytic lineage cancer cells. We observed that the cells start to increase the level of the stemness marker CD133 upon their exposure to α-SYN protofibrils in U-87 MG cells (Figure 3A). TNT biogenesis and cell-to-cell transfer of mitochondria may play a role in mitigating the effects of ROS and rescuing the cells from α-SYN-induced oxidative stress, and the rescued cells proliferate at later time points (12 and 24 hours) (Figure 3B). However, cells start to show a significant increase of the cancer stemness marker CD133 upon treatment with α-SYN protofibrils (Figure 3A-B). We observed increased stemness in U-87 MG cells upon treatment with α-SYN protofibrils revealed by localization of the stem-like marker Nanog in the nucleus maximum at 3 hours (Figure 3C,D). Using WB, we validated increased levels of Nanog (3 to 24 hours) and phospho-Nanog (3 to 24 hours) in the nuclear fraction upon α-SYN protofibrils treatment (Figure 3E,F). Similar results were also observed with U251 cell line (Figure 3G-H).

**Figure 3:**
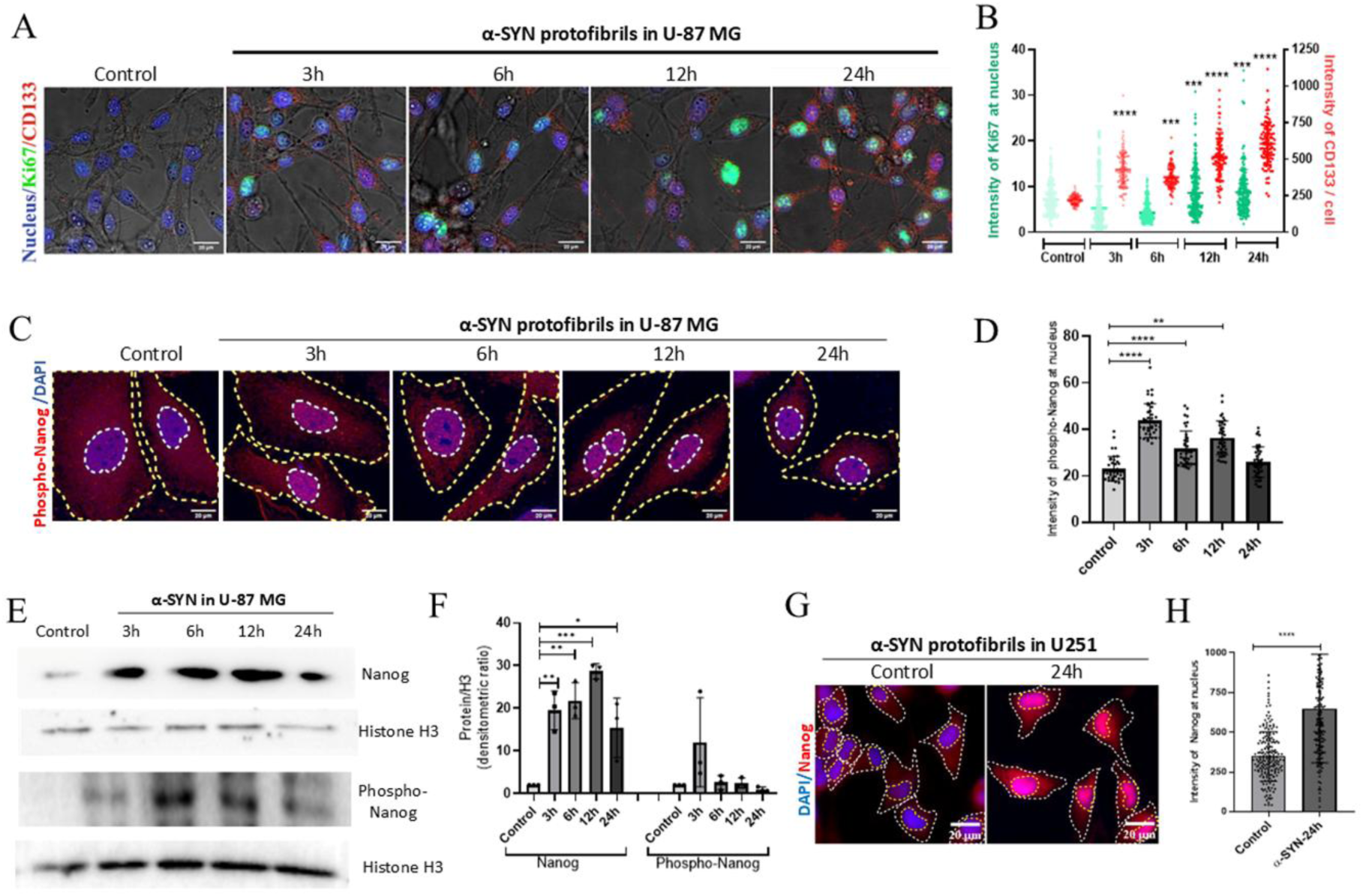
α-SYN protofibrils induce stemness in astroglia. A) Fluorescence images of U-87 MG cells treated with 1µM α-SYN protofibrils with time were stained with proliferation marker Ki67 (green) along with nuclear stain DAPI (blue) and CD133 (red). B) Expressions of Ki67 in the nucleus and CD133 (per cell) were quantified by intensity analysis of the images from the above experiments. C) Fluorescence images of U87-MG treated with α-SYN protofibrils for 24 h were stained with Nanog (red) and DAPI (blue). D) Quantification of nuclear Nanog-positive cells in U87-MG. E) WB of Nanog and phospho-Nanog in the nuclear fraction of α-SYN protofibrils-treated cells. F) Quantification of Nanog and phospho-Nanog from WB data. Full blots of the represented WB bands are shown in Figure S5. G,H) Similarly, Nanog (red) and DAPI (blue) immunostaining in U251 cells and quantification graph. The data points in the graphs show the number of cells included in the intensity analysis, pooling >2 images each from three independent experiments. Data are expressed as mean ± SD, *** p ≤ 0.001. Statistics were analysed using two-way ANOVA. N=3.

### pFAK and Nanog signaling and stemness in α-SYN protofibrils-treated primary astrocytes

Oxidative stress mediated by α-synuclein protofibrils causes the nuclear translocation of phosphorylated FAK, which regulates ROCK-dependent actin remodulation and the biogenesis of TNTs [12]. A report has shown that the nuclear pFAK and Nanog signaling axis may regulate cancer stem cells [41]. Since stemness marker Nanog was upregulated in α-SYN stressed astrocytoma cells, we asked if there is a link between the expression of Nanog and translocation of pFAK in nucleus in astrocytes. To this end, we evaluated the level of pFAK and Nanog in primary astrocytes (Figure 4A) and observed transient nuclear localization of pFAK at earlier time points (3 and 6 hours) and increased nuclear expression of Nanog upon treatment with α-SYN protofibrils during early time-points (Figure 4B,C). Further, we observed co-localization of pFAK (green) with Nanog (red) in the nucleus (3 to 24 hours), which was shown by z-2D and z-stack imaging (Figure 4D). Maximum co-localization was detected at 3 hours (Figure 4E). The findings indicate that nuclear pFAK interacts with the transcription factor Nanog, likely triggering the expression of its downstream target genes.

**Figure 4.**
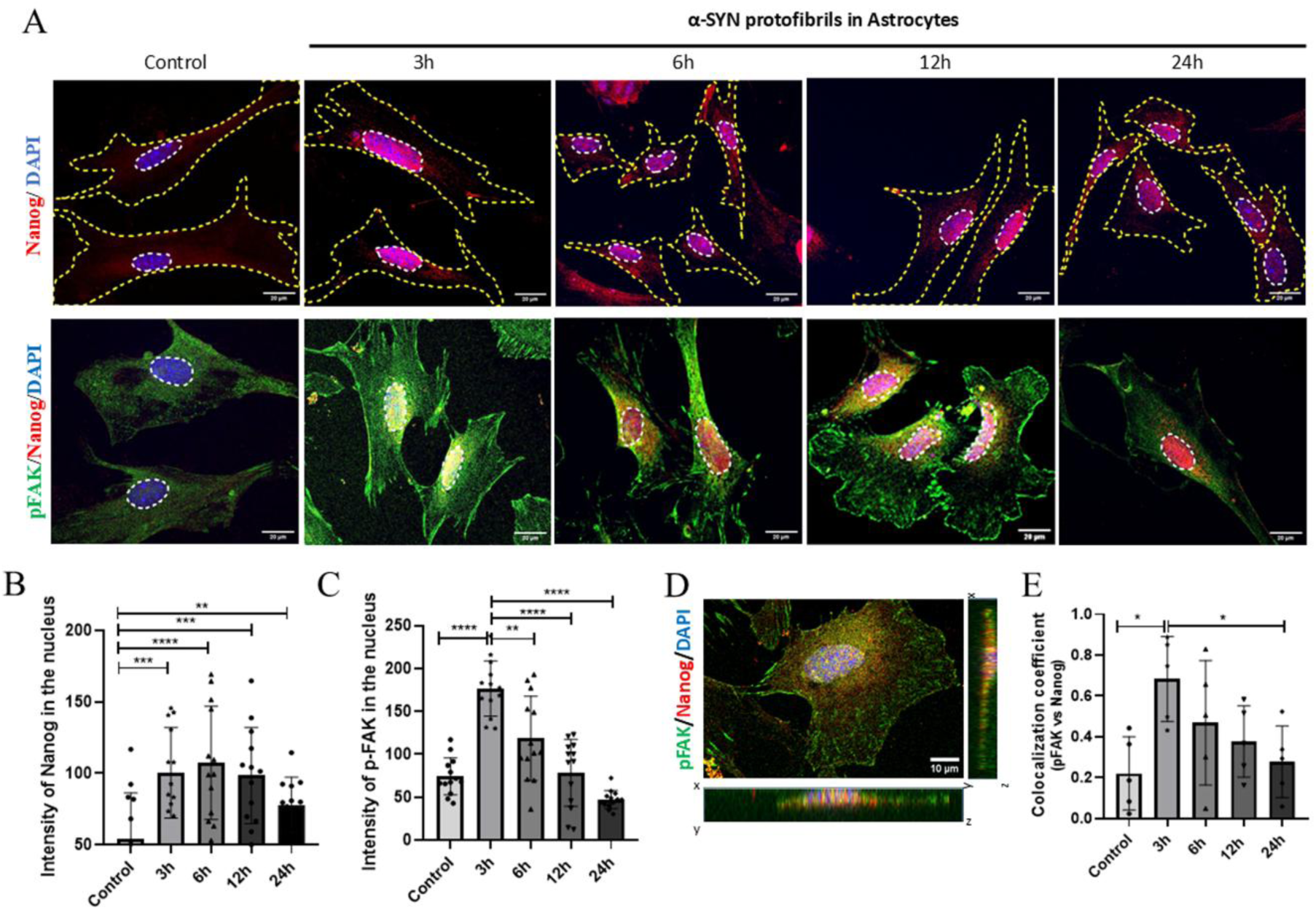
pFAK-Nanog axis in primary astrocytes. Primary astrocytes were treated with 1µM α-SYN protofibrils with time. A) Primary astrocytes were stained with Nanog (red) and DAPI (blue) to detect nuclear expression of Nanog. Double immunostaining was performed to observe expression of Nanog with translocation of nuclear pFAK in primary astrocytes. B) Nuclear expressions of Nanog and C) pFAK in the astrocytes were quantified measuring the intensities. The data points in the graphs show the number of cells included in the intensity analysis, pooling >2 images each from three independent experiments. D) Colocalization of pFAK (green) with Nanog (red) in the nucleus, confirmed through 3D xyz-image plane views. E) Quantification of colocalization coefficient of pFAK vs Nanog in the nucleus of primary astrocytes (analysed from 5 images). Data are expressed as mean ± SD, *** p ≤ 0.001. Statistics were analysed using two-way ANOVA. N=3.

### ROCK-mediated TNT-biogenesis promotes stemness in astroglia

We have shown that α-SYN protofibrils induce increased ROS levels, causing nuclear translocation of pFAK, which regulates the ROCK2 inhibitory signaling pathway and, as a result, promotes formation of TNTs [12]. Interestingly, ROCK inhibitors Y-27632 and fasudil enhance stem-like features of astrocytoma/glioblastoma origin cells [28]. We investigated whether the ROCK inhibitor Y-27632 influences pathways involved in TNT biogenesis to enhance stemness markers in astrocytes. The ROCK inhibitor Y-27632 promoted a significant increase in TNTs in primary astrocytes (Figure 5A), similar to those shown in astroglia cells in our previous study [12]. In addition, treatment with Y-27632 also caused a significant increase in stemness markers Nanog (Figure 5B, E) and SOX2 (Figure 5C,F) in nuclei of primary astrocytes. These findings indicate that inhibiting the ROCK pathway leads to the translocation of pFAK to the nucleus after exposure of cells to α-SYN protofibrils early hours (3 and 6 hours) (Figure 5D), together with increased Nanog levels (Figure 5B).

**Figure 5:**
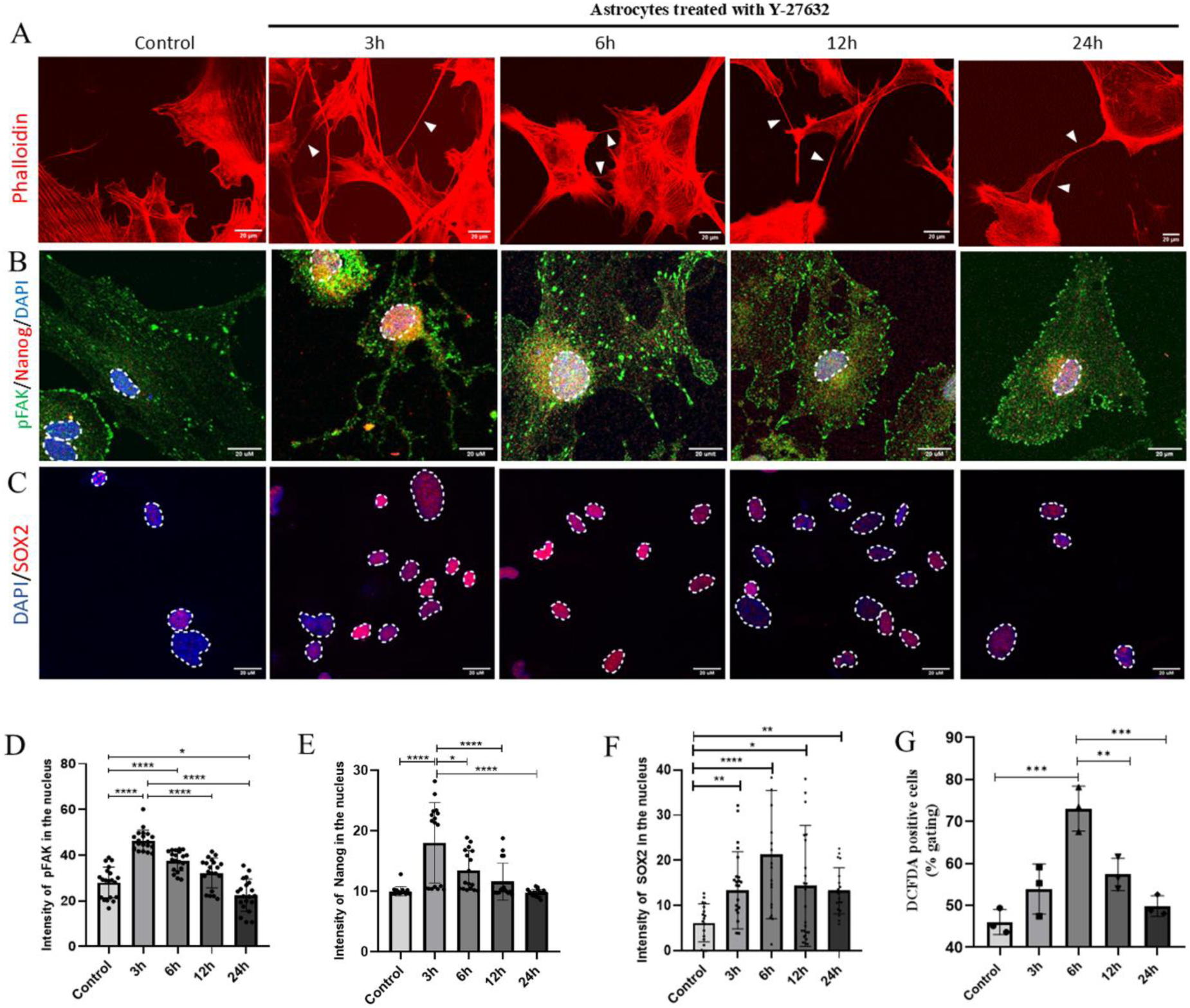
ROCK inhibitor (Y-27632) induces stemness in astroglia. A) ROCK inhibitor (Y-27632) induced biogenesis of TNTs with time in U87 MG cells. Y-27632 treatment causes steady and significant enhancement in stem-cell markes B) Nanog in the nucleus of the primary astrocytes, pFAK colocalizes with Nanog in the nucleus and C) SOX2. Quantification of D) pFAK, E) Nanog, and F) SOX2 intensities in the nucleus of the Y-27632-treated cells. The data points in the graphs represent the mean intensity values of cells from images (2 images from each repeat) pooled across each independent experiment (n=3). G) ROS levels quantified using DCFDA in the flow cytometer in the Y-27632-treated cells, plotted as % gated DCFDA-stained positive cells (data points represent mean values of 3 independent experiments). Data are expressed as mean ± SD, *** p ≤ 0.001. Statistics were analysed using two-way ANOVA. N=3.

The elevated level of nuclear pFAK is transient during the early period after cell exposure, while the increase of Nanog in the nucleus was observed in the ROCK inhibitor-treated astrocytes for a longer time. These results suggest that pFAK-mediated upregulation of Nanog is relatively fast following α-SYN treatment and that the stemness state lasts for extended time periods. Using a co-culture of MitoDsRed (red) and EGFP-life (green) transfected cells, we observed that inhibition of ROCK promoted TNT formation to facilitate transfer of mitochondria in U87 MG cells (Figure S2A-B), ameliorating the level of ROS that increased early following exposure of the cells to the ROCK inhibitor (Figure S2C and Figure 5G). Transfer of mitochondria was significantly more pronounced at 6 hour, when the treatment induced increased numbers of TNTs. Further, we observed the ROCK inhibitor Y-27632-mediated nuclear translocation of pFAK and subsequent increase in the expression of Sox2 and Nanog in the nucleus of U87-MG and U251 cells (Figure S3A-F).

### pFAK-Nanog axis promotes survival of astroglia in α-SYN toxicity by modulating p53 degradation

The RNA-seq differential expression analysis uncovered oxidative stress response genes, several of which are linked to p53-dependent apoptosis. These genes were notably upregulated three hours after treatment with α-SYN protofibrils. However, after 24 hours, the same set of gene expressions was reduced (Figure 6A). We then examined the regulatory mechanisms responsible for the degradation of p53. Our investigation identified a group of genes associated with the MDM2-p53 axis that were upregulated after 3 hours of exposure to α-SYN protofibrils. However, the levels of expression of these genes decreased after 24 hours compared to 3 hours (Figure 6B). MDM2 functions as an E3 ligase that promotes the ubiquitination of p53, leading to its degradation [42]. The signaling pathway involving pFAK and Nanog interacts with p53, leading to its degradation through the E3 ubiquitin ligase Mdm2 [43]. We disrupted the pFAK-Nanog signaling axis by using shRNA to knockdown Nanog and examined the effects of α-SYN protofibrils on p53 during oxidative stress (Figure 6C). We observed an increased expression of p53 in both the cytosol and nucleus at the early time points of 3 and 6 hours after exposure to α-SYN in both U87-MG cells and U87-MG cells transfected with the pLOK.1 vector. The images presented are from wild type (WT) U87-MG cells (Figure 6D-E). However, the levels of p53 significantly decreased at the later time points of 12 and 24 hours. Similar p53 fluctuations with α-SYN treatment were observed in the primary astrocytes (Figure 6F-G). In contrast, exposure of shNanog knockdown cells (with >80% reduced expression, Figure 6H), to α-SYN resulted in significantly higher levels of nuclear p53 during the entire time period (Figure 6D-E). The WB results validate the similar fluctuations in p53 and p21 expression upon α-SYN exposure in control cells (U87-MG and pLKO.1 cells), while p53 and p21 are consistently highly expressed in the Nanog knockdown cells (Figure 6I-K). Quantifications of WBs of p53 and p21 were represented in Figure 6J-K.

**Figure 6:**
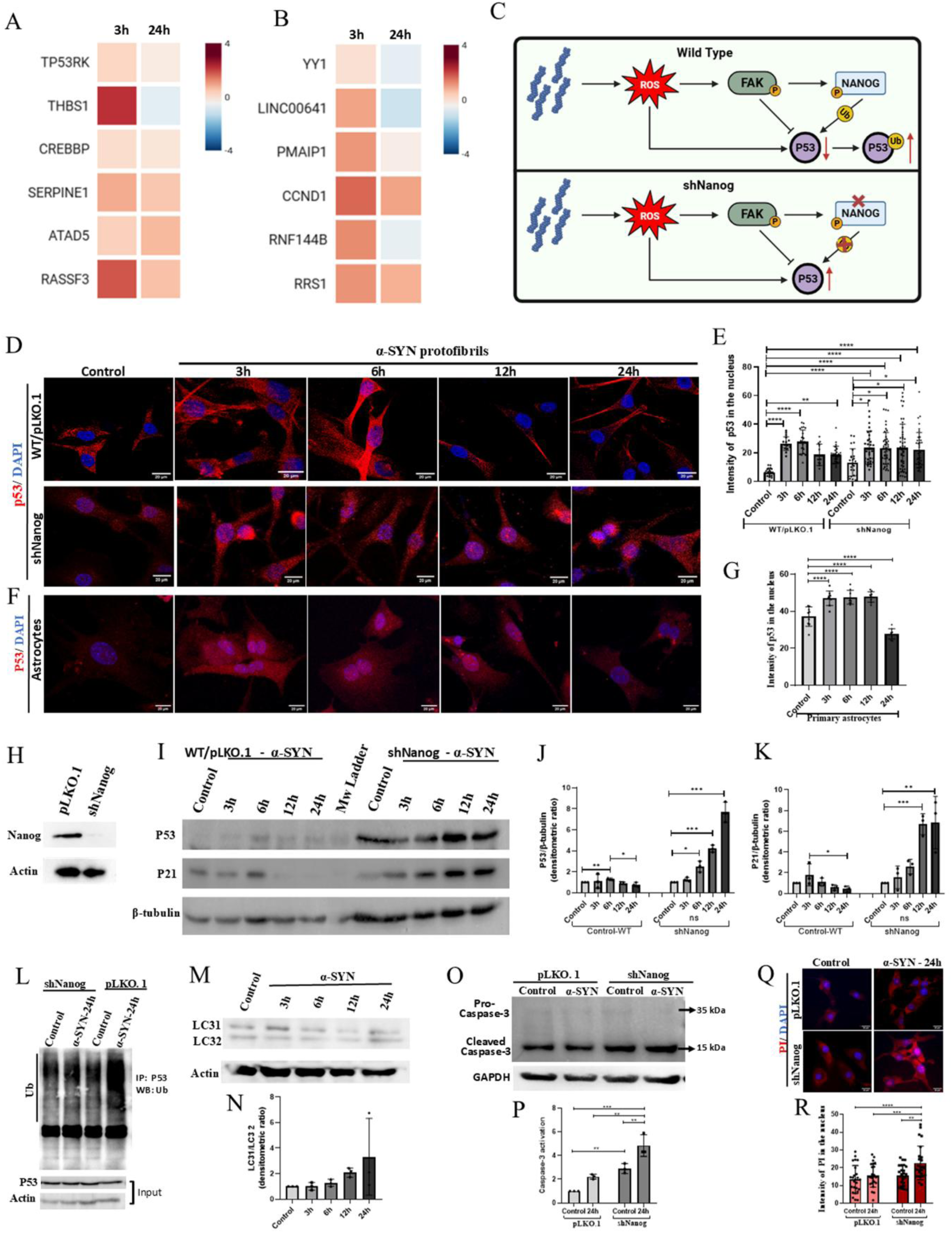
Astroglia survival in α-SYN toxicity by modulating p53 degradation. A) Differentially expressed RNA sequence data were plotted, investigating the genes associated with the oxidative stress response, and linked to p53-directed apoptosis, B) Group of genes associated with the MDM2-p53 axis after 3 and 24 of exposure to α-SYN protofibrils. Data with p-value < 0.05 and adjusted p- value <0.1 were plotted. C) Schematic diagram of pFAK-Nanog signaling axis in α-SYN toxicity-induced p53 modulation by facilitating its degradation. D)Expression of p53 after exposure to α-SYN in U87-MG cell and shNanog knockdown U87-MG cells. E) Quantification of P53 expression in the nucleus of the cells. The data points in the graphs represent the number of cells from images (>6 from each repeat) pooled across each independent experiment (n=3). F) Expression of p53 after exposure to α-SYN in Astrocytes. G) Quantification of P53 expression in the nucleus of the Astrocytes. The data points in the graphs represent the number of cells from images of one repeat. H) The WB of Nanog in pLKO.1 cells and shNanog knockdown cells, with loading control actin. I) The WB results of p53 and p21 expression over time with α-SYN exposure in control U87-MG cells and shNanog knockdown cells. J) Quantification of p53 and K)p21 normalized to loading control β-Tubulin from WB bands. L) IP pulldown of Ubiquitinated (Ub)-p53 in pLKO.1 and shNanog knockdown cells α-SYN treated for 24h with input P53 and loading control Actin. M) The WB of LC3B with α-SYN exposure in U87-MG cells with time, and N) the Quantification of LC31/LC32. O) WB shows pro- and cleaved- caspase-3 to detect apoptosis in pLKO.1 control of U87-MG cells and knocked down shNanog cells treated with α-SYN for 24 h. P) Quantification shows Caspase-3 activation. Q) PI assay to detect apoptosis in U87-MG controls and knocked down shNanog cells treated with α-SYN for 24 h. R) Quantification of PI in the nucleus. Full blots of the represented WB bands are shown in Figure S6 and S7. The data points in the graphs represent the number of cells from images (>6 from each repeat) pooled across each independent experiment (n=3). Data are expressed as mean ± SD, *** p ≤ 0.001. Statistics were analysed using two-way ANOVA. IP experiment N=2 and all other experiments are N=3.

Ubiquitinated p53, which is destined for degradation, preferentially localizes to the cytosol. To assess whether the decline in p53 levels at later time points is a result of its ubiquitin-mediated degradation, we performed immunoprecipitation (IP) pulldown of p53. Increased ubiquitination of p53 is evident in control pLKO.1 cells treated with α-SYN compared to shNanog knockdown cells (Figure 6L). Research indicates that Nanog enhances autophagy during oxidative stress by transcriptionally inducing LC3B (MAP1LC3B), a key protein involved in autophagosome formation [44]. In our RNA sequence (Figure 2), we observed that α-SYN-induced oxidative stress caused a significant downregulation of MAP1LC3B and the autophagy pathway at 3 hours, while its level normalized after 24 hours (Figure 2). The WB data for LC3B indicate increased autophagy at 12 and 24 hours compared to 3 and 6 hours (Figure 6M-N). Degradation of p53 and upregulation of autophagy help protect cells from the proteotoxicity and apoptosis induced by α-SYN protofibrils. We conducted WBs to quantify Caspase-3 activity and performed propidium iodide (PI) staining to detect apoptosis, and our results indicate that U87-MG astroglia cells are resilient to p53-mediated apoptosis by upregulating Nanog through the nuclear translocation of pFAK and the formation of TNT. In cells with knocked-down Nanog, p53 degradation does not occur, and cells promote Caspase-3 cleavage (Figure 6O). Quantification of WBs reveals a significantly higher amount of cleaved Caspase (at 15 kDa Mw) in the shNanog U87-MG cells after 24 hours of α-SYN treatment (Figure 6P). In cells with knocked down Nanog, p53 degradation does not occur, allowing PI to enter these apoptotic cells (Figure 6Q). Quantification reveals a significantly higher amount of PI in the nuclei of shNanog U87-MG cells after 24 hours of α-SYN treatment (Figure 6R).

### Disruption of TNT impairs pFAK-Nanog nuclear co-localization, leading to α-SYN-driven apoptosis

We observed, inhibition of TNTs prevents astroglia cells from surviving and adapting to increased stemness when treated with α-SYN protofibrils. Cyto-D is a known actin depolymerizing agent that inhibits TNTs. Our results show that Cyto-D and ROS inhibitor NAC inhibit α-SYN-induced transient TNTs at early times (3 and 6 hours) in U87-MG cells (Figure 7A). DiD-stained TNTs were counted to quantify the TNT numbers (Figure 7B). Cyto-D and NAC disrupt α-SYN protofibril-induced TNTs, inhibiting the nuclear translocation of pFAK (Figure 7C,F) and preventing the upregulation of Nanog (Figure 7D,G), a key transcription factor for maintaining stemness. Preventing Nanog upregulation stops ubiquitination-mediated p53 degradation in Cyto D treated cells (Figure 7E,H). Cyto-D inhibits actin-cytoskeleton polymerization (Figure 7I), preventing actin modulation that directs pFAK translocation to the nucleus, where NAC partially inhibits ROS, which overall reduces proteotoxic stress. Thus, NAC mediated p53 reduction may happen through alternative pathway (Figure 7E,H). Conversely, the ROCK inhibitor Y-27632 facilitates TNTs via actin remodeling, which in turn promotes the translocation of pFAK to the nucleus and increases the expression of Nanog (Figure 5). To investigate the role of Cyto-D and Y-27632 in α-SYN-induced apoptosis, we conducted Caspase-3 activity experiment (Figure 7J-K) and PI internalization assay (Figure 7L-M). The results indicate that Cyto-D promotes Caspase-3 activation, quantified by the ratio of cleaved- vs pro- Caspase-3. Cyto-D treatment does not prevent apoptosis by inhibiting the biogenesis of TNTs, whereas Y-27632 is effective in preventing apoptosis compared to CytoD by facilitating the formation of TNTs in response to α-SYN protofibril-induced proteotoxicity (Figure 7J-M). These results indicate that inhibiting TNT structures hinders the adaptation of U87 MG astrocytoma cells to stem-like characteristics, stops p53 degradation, and their survival from apoptosis.

**Figure 7:**
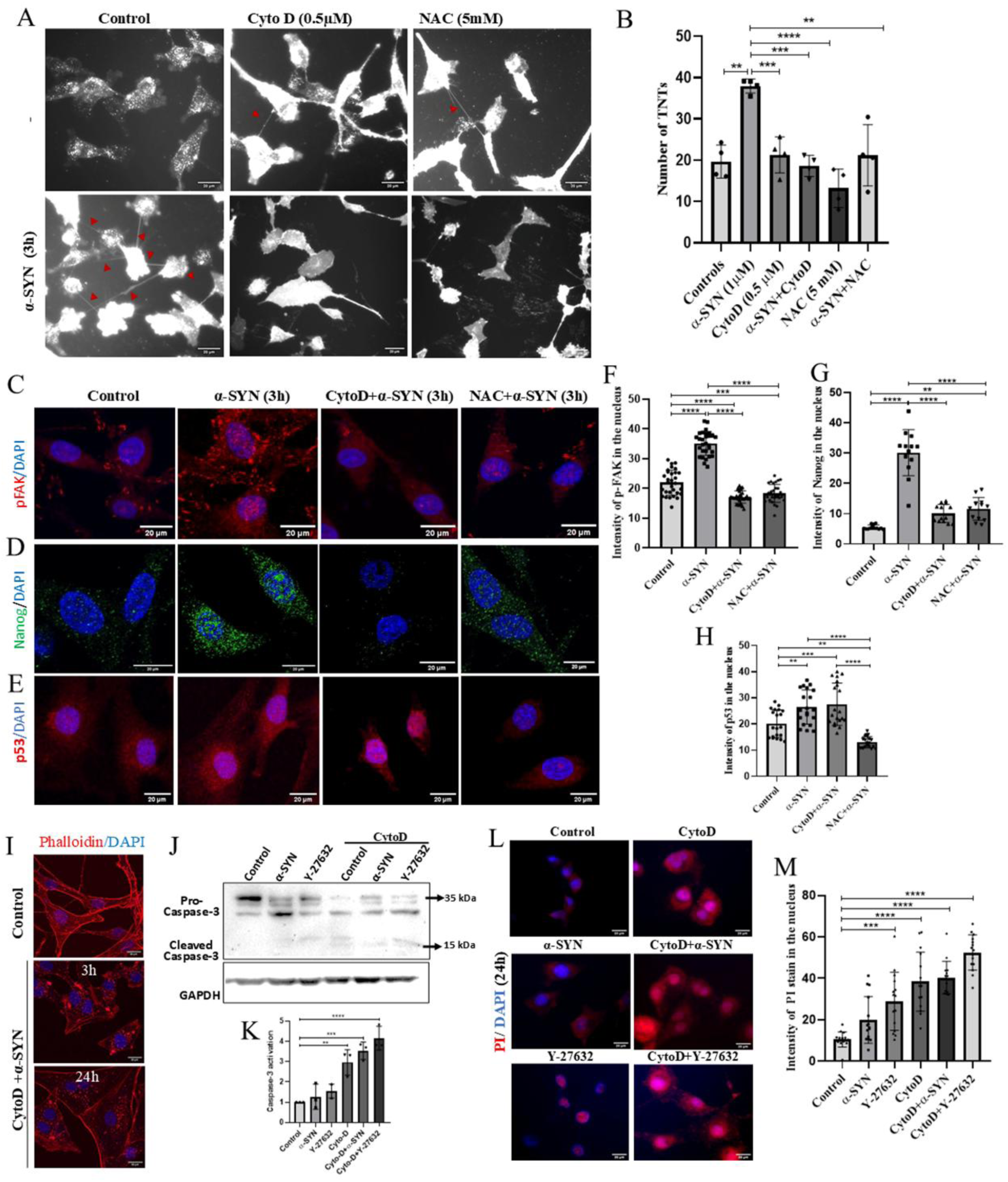
TNT disruption impairs nuclear pFAK-Nanog interaction, triggering α-SYN apoptosis. A) Cyto-D and NAC treatment inhibit α-SYN-induced transient TNTs at early times (3 and 6 h) in U87-MG cells. B) DiD-stained TNTs were counted to quantify inhibition by Cyto-D and NAC. The data points in the graphs represent the average number of TNTs from images of a single representative set. Represented images of nuclear translocation of C) pFAK, D) Nanog. and E) p53 with Cyto-D and NAC treatment. Quantification of nuclear F) pFAK, G) Nanog and H) p53. The data points in the graphs represent the number of cells from images (>4 per repeat) pooled across independent experiments (n=3). I) Cyto-D inhibits actin-cytoskeleton polymerization. J) WB to detect Caspase-3 activity and L) PI internalization assay in the presence of Cyto-D and Y-27632 in α-SYN-treated cells. Quantification of K) Caspase-3 activation and M) PI internalization. Full WBs were shown in Figure S7. The data points in the graphs represent the number of cells from images (>4 per repeat) pooled across independent experiments (n=3). Data are expressed as mean ± SD, *** p ≤ 0.001. Statistics were analysed using two-way ANOVA. N=3.

### Validation of increased stemness after treatment with α-SYN protofibrils and ROCK inhibition

We used the hanging drop 3D-spheroid assay in order to assess the stemness state in astroglia treated for 24 hours with α-SYN protofibrils and the ROCK inhibitor Y-27632. Our results show significantly greater sizes of spheroids upon treatment with α-SYN in primary astrocytes compared to control cells (Figure 8A-B). Larger sizes of spheroids were observed upon treatment with α-SYN and the ROCK inhibitor separately and in combination in the U251 and U87-MG cells (Figure 8C). Treatment with the actin-depolymerizing agent Cyto-D, a general inhibitor of TNTs, prevented spheroid formation (Figure 8C). The size of spheroids reached a maximum at 6 hour after α-SYN treatment, and consistent spheroid formation with significantly larger sizes persisted for up to 24 hours in the treated cells (Figure 8D). The ROCK inhibitor also led to a significant increase in spheroid size. Combination of the inhibitor with α-SYN causes a further increase in the spheroid size, which peaked 6 hour after treatment, suggesting a synergistic effect (Figure 8D). This observation suggests that α-SYN protofibrils likely inhibit the ROCK signaling pathway to enhance the biogenesis of TNTs, which leads to an increased level of stemness. We further observed that inhibition of spheroid formation occurred upon Nanog knockdown in U87-MG cells following treatment with α-SYN protofibrils (Figure S4). This suggests that the interaction of pFAK with the transcription factor Nanog plays an important role in changes at the transcription level, leading to increased stemness. U87 MG and U251 astrocytoma/glioblastoma cells contain a large population of cancer stem-like cells, which increases following α-SYN treatment. Notably, primary astrocytes also develop a substantial pool of stem-like cells after exposure to α-SYN protofibrils, potentially contributing to enhanced proliferation.

**Figure 8:**
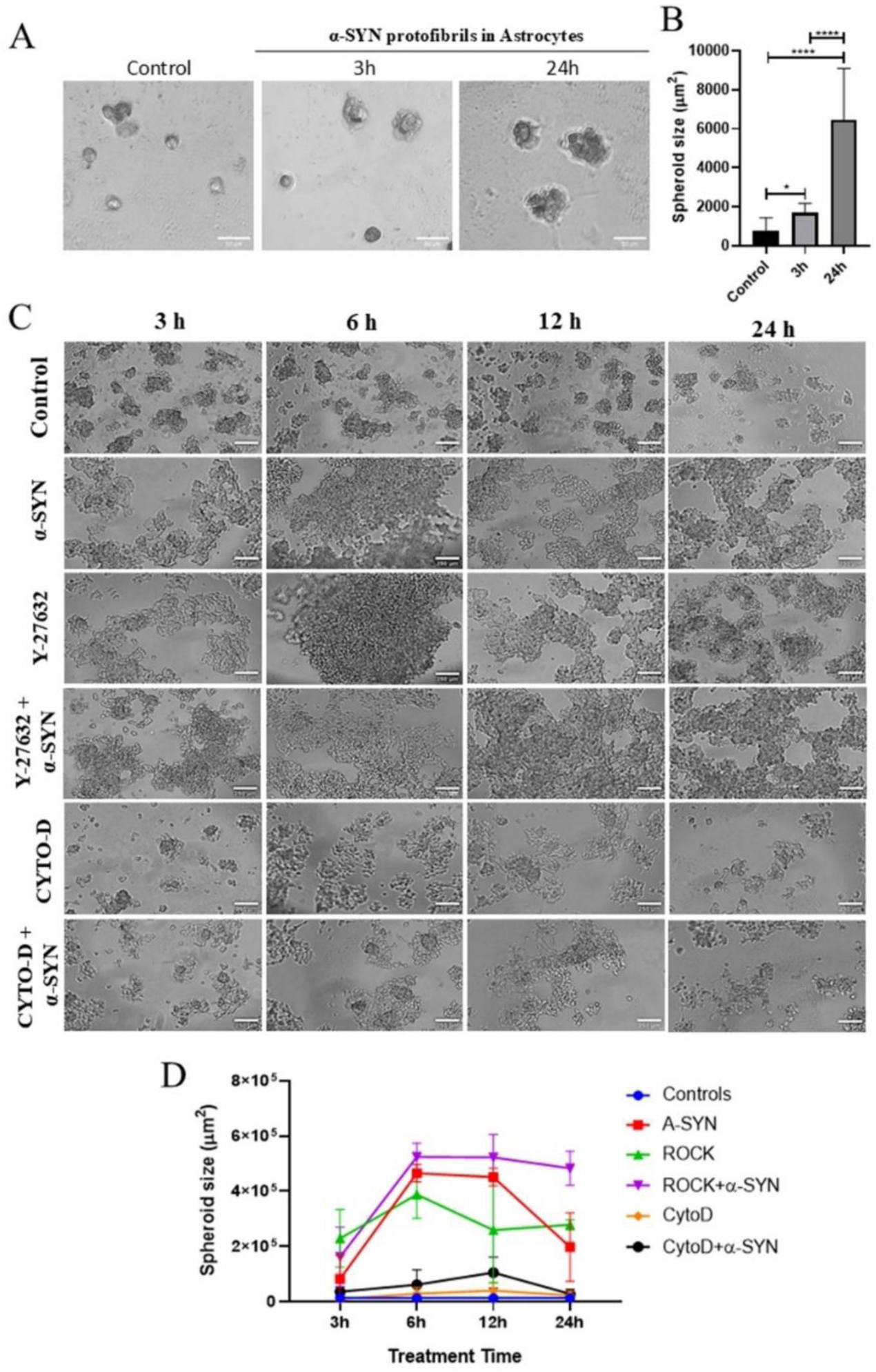
Hanging drop spheroid assay to assess stemness. **A)** The panel represents the growth of spheroids of primary astrocytes upon treatment with α-SYN (1 μM) after 3 h and 24 h of treatment. B) The graph represents the size of spheroids of the treated primary astrocytes. C) The panel represents the growth of U251 spheroids upon treatment with α-SYN (1 μM), ROCK inhibitor (Y-27632, 5 μM), Cyto-D and α-SYN + Y-27632 for 3h, 6h, 12h and 24h. D) The graph represents the size of spheroids of the treated U251 cells. The quantification graphs shown for spheroids are from 6 image fields of view in a set (n=3). Data are expressed as mean ± SD, *** p ≤ 0.001. Statistics were analysed using two-way ANOVA. N=3.

## Discussion

Under normal physiological conditions, astrocytes function as supportive glial cells within the CNS. They modulate neurotransmission, respond to neurodegeneration, neuroinflammation, and injury. Studies indicate that astrocytes can serve as precursors of gliomas, similar to neural stem cells and oligodendrocytes. Oncogenic mutations introduced via GFAP-CreER in the adult brain are the major cause for over 20% of tumours arising from astrocytes, apart from those arising from neural stem cell niches [45]. Additionally, the tumor microenvironment and its communication with surrounding healthy astrocytes promote tumour progression by shaping metabolic interactions. It has been reported that the spread of α-SYN from glioblastoma cells to adjacent astrocytes via TNTs results in the development of stem-like properties [24]. TNTs play a crucial role in the rescue of cells from deleterious effects of toxic α-SYN by facilitating cell-to-cell transfer of mitochondria in neuroglial cells [19]. Additionally, TNT-mediated intercellular transfer of mitochondria from glioblastoma cells to surrounding astrocytes leads to the adaptation of healthy cells to metabolically aberrant tumour cells with stem-like properties [23]. This process is complex and involves various molecular changes that lead to the transformation of astrocytes into cancer stem-like cells.

There is an emerging role of TNTs in protecting astrocytes and astrocytic lineage cells against stress induced by α-SYN protofibrils. This study illustrates that proteotoxic stress from α-SYN leads to mitochondrial damage, oxidative stress, and elevated levels of ROS, resulting in a significant increase in mitochondrial membrane potential (ΔΨm), creating a cytotoxic environment that threatens cellular homeostasis. Consequently, this process facilitates the formation of TNTs via modulation of ROCK signaling via nuclear translocation of pFAK [12]. Notably, upon α-SYN challenge, pFAK relocates to the nucleus of astrocytes and astrocytoma cells, where it activates Nanog and colocalizes with this transcription factor, which is traditionally involved in maintaining pluripotency in stem cells. The interaction between pFAK and Nanog within the nucleus appears to be a pivotal event in the α-SYN-treated astrocyte’s adaptive mechanism to protect cells from oxidative stress-induced apoptosis.

TNTs are nanotubular structures serving as conduits for the exchange of not only pathogenic protein aggregates but also damaged and functional mitochondria. Importantly, formation of TNTs is not a static or constitutive process; it is tightly regulated, transient, and activated specifically in response to mitochondrial stress signals that are initiated by α-SYN. It was also observed that the cells switch to a glycolysis-dependent metabolic pathway to meet their energy needs, as functionally aberrant mitochondria cannot sustain energy requirements via oxidative phosphorylation. This mechanism is similar to the Warburg effect in cancer [36]. These changes promote faster cell growth and expansion, as well as trigger pro-survival pathways [48]. Studies show that TNTs can mediate transfer of healthy as well as aberrant mitochondria, which restores energy metabolism and promotes cell proliferation, respectively. The underlying mechanism driving these conflicting actions remains elusive [12], [22], [46], [47].

RNA sequencing analysis in this study also indicates a metabolic shift from oxidative phosphorylation to glycolysis, and reduced mitophagy with transient TNT formation at early time points while mitophagy is enhanced at later time points, attributed to TNT-mediated ROS mitigation and cellular adaptation. These data suggest that TNTs play a primary role in reducing ROS caused by aberrant dysfunctional mitochondria during early stages of α-SYN treatment, surpassing mitophagy in this function. A recent study demonstrates that astrocytes activate autophagic flux to eliminate dysfunctional mitochondria and support cell viability after internalizing extracellular α-synuclein [49].

One of our intriguing observations is that modulation of the tumor suppressor protein p53 during TNT formation upon α-SYN response. P53 is a key protein that plays a central role in apoptosis and DNA damage responses. Signaling axis of pFAK and Nanog crosstalk with p53 facilitates its degradation via the E3 ubiquitin ligase Mdm2 [43]. This process not only prevents p53-mediated apoptosis but also reprograms the cell toward a more resilient, stem-like state. Altogether, these findings suggest that under proteotoxic stress, astrocytes temporarily adopt a stem-like phenotype that enhances their survival and protective functions. This cellular remodeling is characterized by upregulation of transcription factors related to stemness and the signaling pathway that facilitates TNT formation.

We show that in α-SYN protofibrils-treated senescence-like cells, pFAK translocated to the nucleus, colocalizes with Nanog, enhancing the stem-like progenitor markers in astroglia cells. Research has demonstrated that the transcription factor Nanog attaches to the FAK promoter, enhancing FAK expression. Additionally, FAK binds to Nanog and phosphorylates it, and both are essential for regulating cancer stem cells [43]. We have also demonstrated that the ROCK inhibitor Y-27623 enhances stemness in astroglia cells by regulating ROS levels and directing pFAK to the nucleus. In this study, we establish using a 3D-spheroid assay that treatment with α-SYN protofibrils and a ROCK inhibitor enhances stem-like markers in terminally differentiated primary astrocytes and as well as astrocytoma cells. There are studies showing that ROCK inhibition is linked to metabolic shifts, including enhanced glycolysis and reduced mitochondrial respiration, which may elevate oxidative stress [50]. A multitude of cancer cells exhibit accumulation of senescent cells as their precursors, which may enhance the invasion of cancer cells in response to aberrant signaling pathways by inducing proliferation and stemness [51].

In contrast, inhibition of key steps in the TNT formation pathway disrupts this protective mechanism, with cells proceeding towards apoptosis. Numerous studies suggest that TNTs may help protect cells from P53-mediated apoptosis and enhance their resistance to various treatments [17], [35]. While it has been unclear how P53 is involved in TNTs, our study demonstrates that P53 is not the cause of TNT formation; instead, the TNT biogenesis signaling pathway modulates P53 degradation. Cyto-D, an actin-depolymerizing agent, effectively prevents the formation of TNTs by disrupting the actin cytoskeleton, which is crucial for its structure. Treatment with NAC prevents the translocation of pFAK to the nucleus and inhibits the formation of TNTs, resulting in disruption of its interaction with Nanog. Both interventions result in an increase in α-SYN-induced apoptosis, suggesting that TNTs are active participants in the cellular survival response rather than mere by-products of proteotoxic stress.

Further confirming the central role of Nanog, knockdown of Nanog results in persistence levels of p53, compromising astroglia protection against apoptosis. In these cells, even when pFAK translocates to the nucleus, the lack of Nanog prevents the necessary downstream signaling for p53 degradation. This highlights another unique function of Nanog in cellular survival mechanisms initiated by pFAK translocation, in addition to its well-established role in embryonic stem cells. It is important to note that this type of cellular reprogramming is temporary and does not indicate a transition to a fully pluripotent state; instead, it highlights a temporary alteration in gene expression that enhances the astrocyte’s ability to survive and provide neuroprotection.

The study explains how the TNT biogenesis pathway helps reduce proteotoxic stress, thereby aiding cellular adaptation. It alleviates mitochondrial stress and oxidative damage, promotes the degradation of P53 to protect cells from apoptosis, and simultaneously enhances the expression of stem-like phenotypes. TNTs mediate the adaptation of astrocytoma/glioblastoma cells to promote their resistance to Tomozolomide and ionization radiation [52] and facilitate non-tumor astrocytes to adapt to metabolic tumor conditions such as hypoxia [23]. Several mutations identified in genes associated with familial cases of PD are notably linked to increased risk of brain tumors [53]. These associations became more noticeable when data were analysed considering temporal basis and certain ethnic groups [54], [55]. Several common genes and mutations associated with aberrant α-SYN aggregation, mitochondrial dysfunction, and cell cycle dysregulation, such as SNCA, PARK2, PARK8, PTEN, MC1R, ATM, and p53, have been implicated in this positive association [54].

## Conclusion

In conclusion, this study reveals that α-SYN protofibrils induce mitochondrial stress and ROS in astroglia, triggering transient nuclear translocation of pFAK to promote transient TNT formation, whereby it co-localizes with Nanog. The interaction between pFAK and Nanog promotes degradation of p53 via Mdm2 and autophagy, which in turn prevents apoptosis. These transient TNTs enable the exchange of mitochondria between cells, helping to restore redox balance and improve the survival of astroglial cells. Inhibition of TNTs or Nanog disrupts the protective mechanisms that prevent apoptosis induced by neurodegenerative proteotoxicity in astrocytes. The study describes how cells facilitate the temporary formation of TNT structures under conditions of metabolic stress or aberrant states, and how the p53 pathway is indirectly connected to the development of transient TNT formation. Overall, pFAK-mediated TNT formation is a critical adaptive response that connects mitochondrial homeostasis, stemness signaling, and cellular resilience in astroglia during apoptosis induced by α-SYN protofibrils-induced proteotoxic stress.

## Authors’ contributions statement

SN and RK conceptualized the work; RK, ASBK, RRM, AK, SP (JNCASR) and SJ performed the experiments; VMR and SGD performed RNA sequencing analysis; RK, ASBK, and SN designed the work; RK, ASBK, SP (NIMHANS), VKR, RM, JN, and SN validated and investigated all the formal analyses and interpretation of data; VKR, SGD, SP (NIMHANS), and RM provided methodology support; RK and SN wrote the first draft; JN provided significant intellectual contribution; All authors edited and commented on the manuscript. All authors read and approved the final manuscript.

## Acknowledgements

We thank Ms Abinaya Raghavan, Ms Suchana Chaterjee and Ms Ahana Das for their contribution to the graphical abstract and for helping with making lab reagents. The authors thank Genotypic Technology for supporting next-generation sequencing and analysis of the data. We thank Prof. Anujith Kumar for providing a few reagents. We thank Ms. B. Suma for the confocal microscopy, the JNCASR confocal facility, and the MIRM-MAHE Bangalore microscope facility. The authors acknowledge Grammarly software for English editing. The authors declare that they have not used AI-generated work in this manuscript.

## Ethics approval and consent to participate

All experimental protocols were approved and conducted in accordance with national policies for the use of animals, regulated by the KMC, Manipal Academy of Higher Education, Manipal, India. (Title of approved project: Tunneling nanotubes in rescuing of neurodegenerative toxic burdens by facilitating glial cross-talk through cell-to-cell transfer and its role in reversal of senescence, approval number: IAEC/KMC/75/2023, date of approval: October 13, 2023). Mice were euthanized by CO₂ asphyxiation followed by cervical dislocation to confirm death, in accordance with institutional animal care and use guidelines. All animal experiments were conducted and reported in accordance with the ARRIVE guidelines 2.0.

This declaration confirms that the original source, ECACC, of human cell lines U87 MG and U251, has verified that initial ethical approval was obtained for the collection of human cells and that the donors provided informed consent.

## Consent for publication

All authors read and approved the final version of the manuscript.

## Data and code availability

Raw data obtained from RNA-seq have been deposited at GEO (GSE295417). Additional data pertaining to this article will be shared upon request to the lead contact. We do not report any original code in this article.

