## Supplementary material for "α-Synuclein Triggers Intercellular Nanotubes Formation to Prevent Apoptosis in Astroglia by Promoting Stemness": Supplemantary Figures

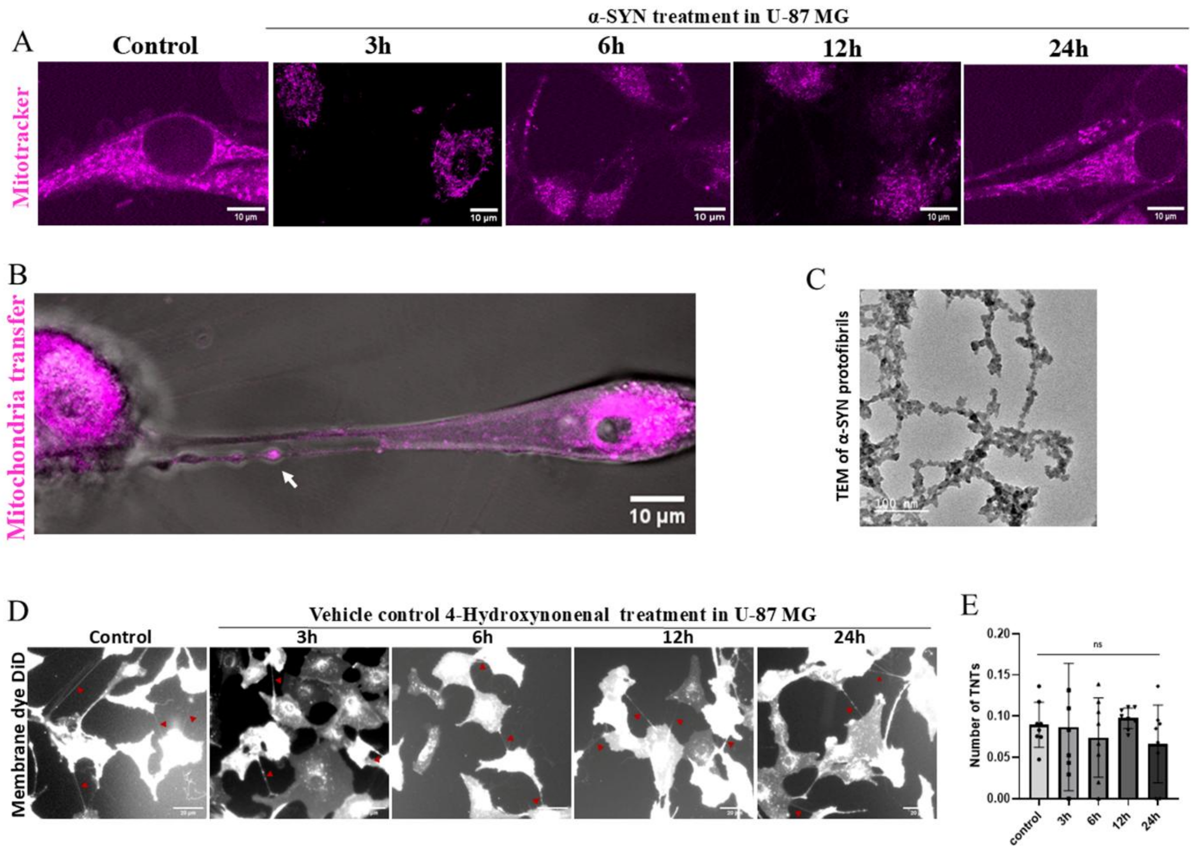


***Figure S1: Effect of α-SYN protofibrils (1 μM) on mitochondrial membrane potential.*** *A) Mitotracker (magenta-red) stained U87-MG cells treated with α-SYN protofibrils (1 μM) for 3, 6, 12, and 24 hours. Images show that early exposure to α-SYN resulted in fragmented mitochondria and reduced intensities of mitotracker stains. B) α-SYN treatment facilitates TNTs at early time points (3h), and mitochondria travel through the TNTs from one cell to another. C) TEM images of α-SYN protofibrils. D) Vehicle control of 4-Hydroxynonenal treatment shows no effect on TNT formation over the treatment duration. E) Quantification of TNT numbers over the treatment duration upon addition of vehicle control media in the U-87 MG cells. Data are expressed as mean ± SD, *** $p \leq 0.001$. Statistics were analysed using two-way ANOVA. N=3.*

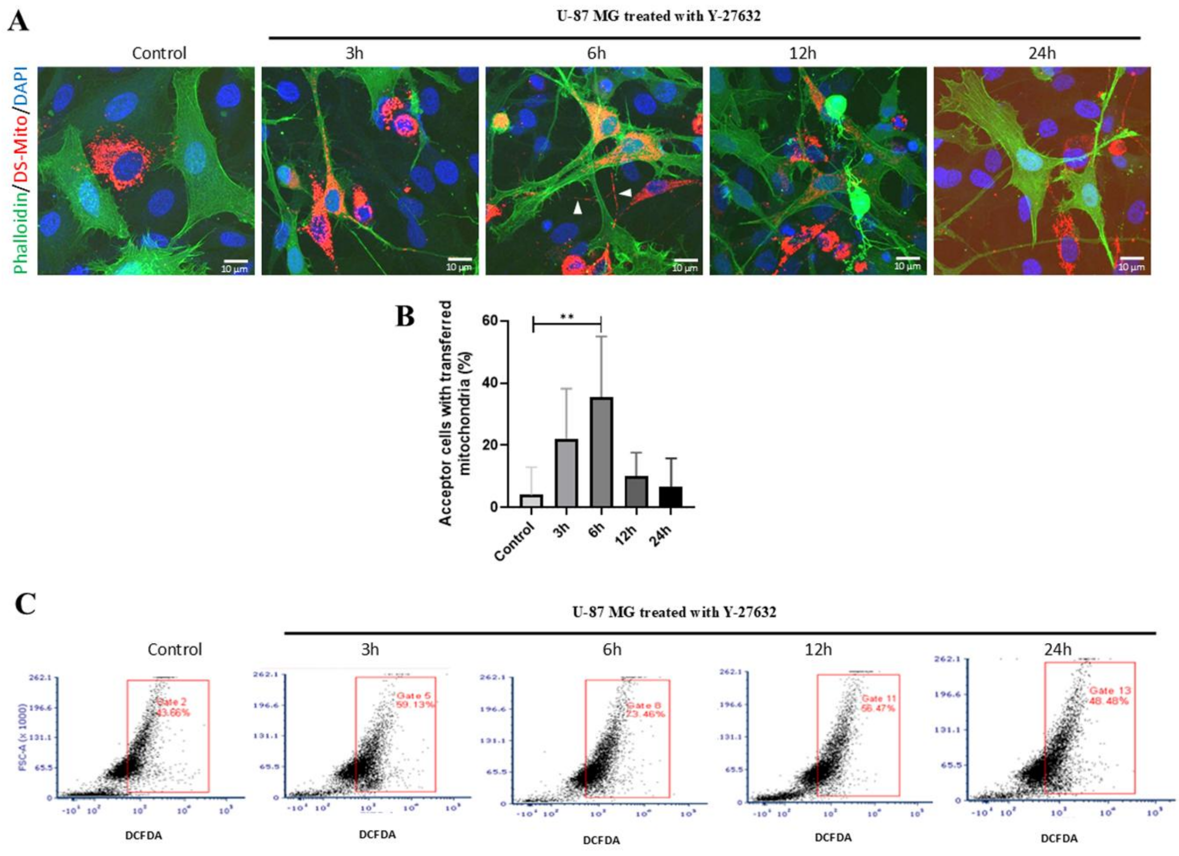


***Figure S2.*** *A) Co-culture with MitoDsRed (red) and EGFP-life (green) transfected cells, shows that ROCK inhibitor (Y-27632) promotes TNT formation and facilitates the transfer of mitochondria (red) in the EGFP-life (green) transfected U87-MG cells. B) Quantification of mitochondria transfer shows that the transfer is maximum at 6h of Y-27632 treatment. The number of cells with transferred mitochondria was counted manually (counted from randomly taken > 10 images for each set). Data are expressed as mean ± SD, *** $p \leq 0.001$. Statistics were analysed using two-way ANOVA. N=3. C) The represented DCFDA positive U87-MG cells were gated to analyse the ROS levels. Results show that the cell-to-cell transfer of mitochondria ameliorates the level of ROS that increase early in the ROCK inhibitor-treated cells. The quantification of the represented panels are demonstrated in the Figure 4G. N=3.*

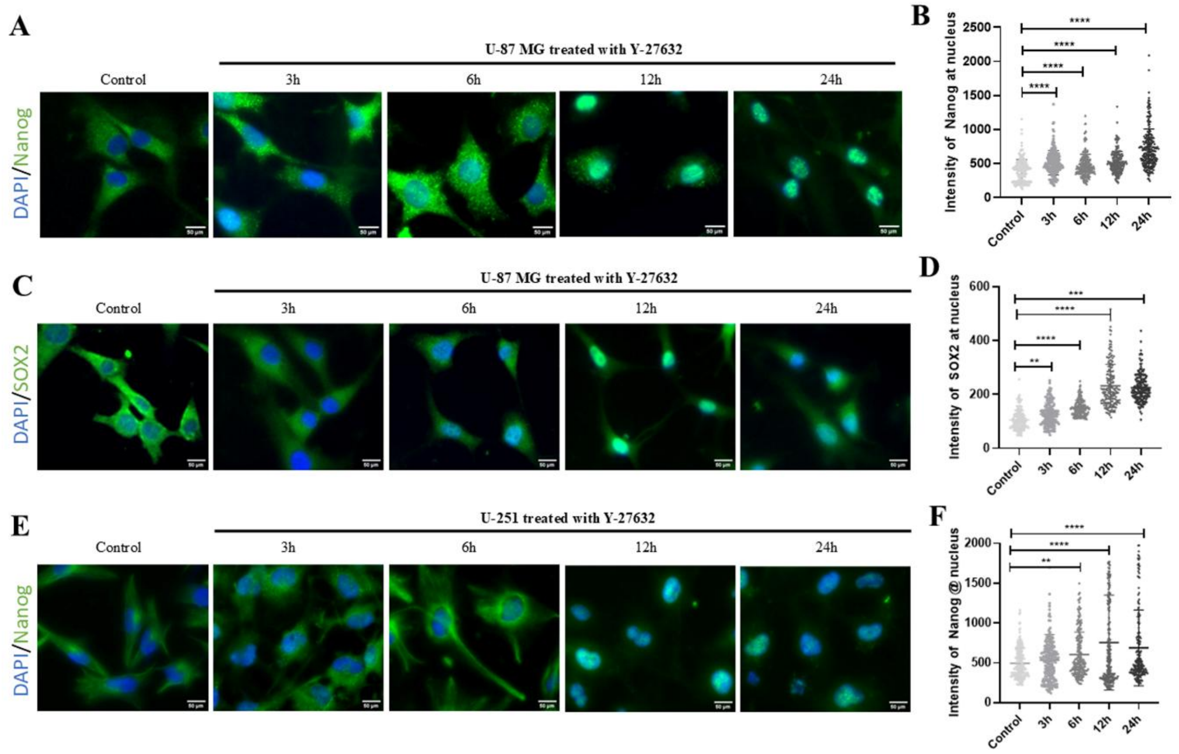


***Figure S3:*** *ROCK inhibitor y-27632 treatment, causes an increase in the expression of Nanog and Sox2 in the nucleus of U87-MG (A,B and C,D) and U251 cells (E,F) over time. Nuclear Nanog and Sox2 positive cells were quantified using Image J. Image analysis was done from > 150 cells for each set. Data are expressed as mean ± SD, *** p ≤ 0.001. Statistics were analysed using two-way ANOVA. N=3.*

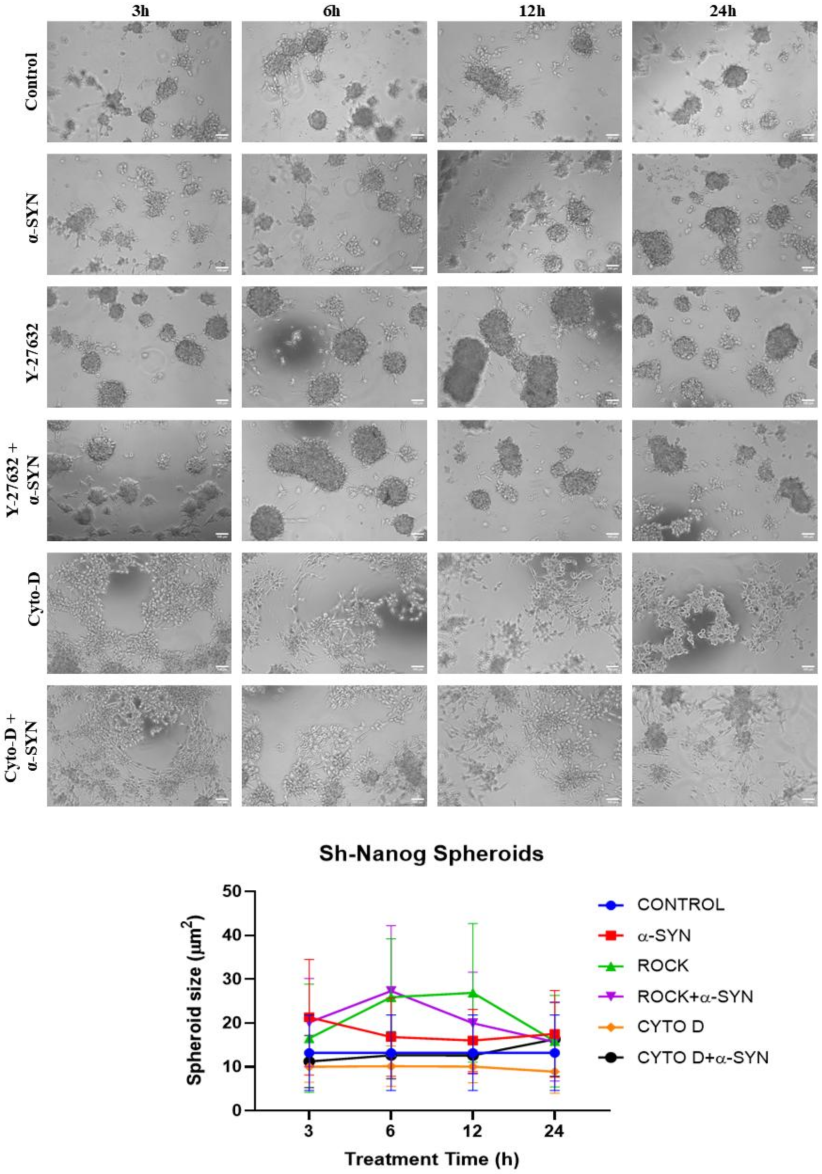


***Figure S4: Hanging drop spheroid assay to evaluate stemness in U87-MG cells with Nanog knockdown via shRNA.*** *The panel represents the prevention of growth of spheroids in the Nanog knockdown U87-MG cells upon treatment with α-SYN (1 μM), ROCK inhibitor (y-27632, 5 μM), cytochalasin-D and α-SYN + y-27632 for 3h, 6h, 12h and 24h, compared to pLKO.1 control cell and normal control cells (Figure 6B). The graph represents the size of spheroids of the treated cells. Data are expressed as mean ± SD, *** $p \leq 0.001$. Statistics were analysed using two-way ANOVA. N=3.*
